# Establishing Essential Oil Stewardship Through the Case of Rosemary and Thyme Oils Against *Staphylococcus aureus*

**DOI:** 10.1101/2025.07.16.665039

**Authors:** Malwina Brożyna, Zuzanna Stępnicka, Katarzyna Kapczyńska, Bartłomiej Dudek, Adam Matkowski, Adam Junka

## Abstract

Essential oils (EOs) have long been studied for their antimicrobial properties, yet most investigations rely on simplistic models, limited strain panels, and anecdotal interpretations - failing to meet the standards expected of modern anti-infective agents. Advancing beyond this tradition, we implement a framework developed over several years of systematic investigation. Using this approach, we assessed the antibiofilm activity of *Rosmarinus officinalis* L. and *Thymus vulgaris* L. EOs against a panel of clinical *Staphylococcus aureus* isolates obtained from non-healing wounds. By applying infection-relevant conditions, such as wound-mimicking media and surfaces, strain-level resolution, and both contact and volatile exposure, we revealed substantial inter-strain variability in susceptibility, challenging the notion of EOs as uniformly effective agents. This variability was quantified using robust statistics, lending confidence to the reproducibility and translational relevance of the findings. These results underscore the need for Essential Oil Stewardship: a reproducible, interdisciplinary framework for EO testing, interpretation, and clinical translation. Our work demonstrates that such approach is feasible and sets the foundation for its broader adoption. The key message from this study is that EOs cannot meaningfully support or complement antibiotics and antiseptic agents in combating infections unless they are evaluated with the same methodological rigor.

## 1 Introduction

Antimicrobial resistance (AMR) is no longer a hypothetical future concern - it is a present and escalating global health crisis. According to the report published in 2022, AMR was directly responsible for an estimated 1.27 million deaths in 2019 and was associated with nearly 5 million more. Forecasts indicate that drug-resistant infections could claim up to 10 million lives annually by 2050, surpassing cancer as the leading cause of death worldwide. Such projections underscore the declining effectiveness of antibiotics and the resulting urgency to develop alternative strategies to support them (Naghavi et al., 2024).

One such strategy has involved the topical antiseptic agents as the defense in the management of localized skin infections, non-healing wounds or mucosal membranes. Their appeal stemmed from several advantages: direct topical application, reduced systemic exposure, and a low likelihood of resistance development due to a broad, rapid mode of action (Punjataewakupt et al., 2019). Compounds such as povidone-iodine, chlorhexidine, octenidine, or polyhexanide became central to clinical protocols across Europe and beyond. However, povidone-iodine, though highly potent, is associated with contraindications in certain patient populations (Eggers, 2019). In turn, chlorhexidine has faced increasing scrutiny due to reports of emerging resistance and hypersensitivity reactions (Kampf, 2016; Opstrup et al., 2019). As chlorhexidine use declined, octenidine emerged as a preferred alternative across both hospital and community settings. However, early evidence of reduced susceptibility in certain pathogens towards this antiseptic has already been reported (Kampf, 2018). This highlights that extensive and sustained antiseptic use imposes evolutionary selective pressure, driving the development of microbial resistance in a manner analogous to what has been documented for antibiotics. Thus, one of the emerging directions in the search for sustainable antimicrobial solutions has been a renewed exploration of natural products, particularly those evolved by plants as part of their own defense systems.

Among these, essential oils (EOs) have attracted considerable interest due to their complex, multicomponent nature and broad biological activity. Unlike conventional (obtained via chemical synthesis) antiseptic products, which typically rely on a single active compound, EOs are composed of dozens of constituents (Swamy et al., 2016). These mixtures do not act solely as antimicrobials; many components exhibit anti-inflammatory, antioxidant, and modulatory properties, offering a multifaceted therapeutic profile (Spisni et al., 2020). Importantly, their antimicrobial mechanisms of action are nonspecific, targeting, among others, bacterial membranes, enzymes, and signaling pathways simultaneously - thereby reducing the likelihood of resistance development. This stands in contrast to the reductionist paradigm that has long dominated XIX and XX-century medicine: one pathogen, one drug, one target (Kon and Rai, 2012).

EOs represent a different antimicrobial strategy. Their chemically diverse composition functions as an evolutionarily selected form of broad-spectrum preparedness, offering concurrent activity against multiple microbial targets without prior pathogen-specific adaptation (Kon and Rai, 2012). Such approach may now offer valuable inspiration for the post-antibiotic era. An additional advantage of EOs lies in their dual-phase activity: they can act both in the liquid and volatile phase (Bunse et al., 2022). This allows EOs to reach microenvironments that are otherwise inaccessible to topically applied liquids, such as the deeper layers of biofilms or poorly perfused wound niches.

Given these appealing properties (broad-spectrum antimicrobial activity, low cytotoxicity, and physicochemical versatility), EOs are often portrayed as ideal candidates for adjunctive or alternative anti-infective therapies (Hyldgaard et al., 2012; Ju et al., 2022). Yet despite the frequent repetition, the methodological quality underpinning these statements remains inconsistent. Many studies rely on low-resolution assays, single reference strains, or simplistic viability readouts that fail to capture the interactions between EO components, microbial physiology, and the complex microenvironment of the infection sites, further complicated by the ongoing host-pathogen interplay.

Such studies often disregard both the physicochemical heterogeneity of EOs and the complexity of their targets. A particularly problematic example is the aromatogram - a crude adaptation of the classical disk diffusion method, in which multiple EOs or different concentrations of the same EO are applied onto the same agar plate (Calvo et al., n.d.; Meroni et al., 2020; Chama et al., 2022). The resulting zones of inhibition are difficult to interpret and virtually impossible to standardize. They do not reflect the activity of any single oil but are shaped by an uncontrolled combination of liquid-phase diffusion and airborne transfer of volatile fractions (Brożyna et al., 2020).

Another limitation in EOs research is the lack of alignment between experimental models and the physiological realities of infection sites. Standard antimicrobial assays often rely on nutrient-rich laboratory media, polystyrene surfaces, and planktonic cell suspensions - i.e., conditions that poorly reflect the landscape of actual infectious niches (Thaarup and Bjarnsholt, 2021).

Non-healing wounds represent a particularly illustrative example of this mismatch. These environments are defined by fluctuating oxygen tension, persistent inflammation, polymicrobial consortia, and dense biofilms embedded within polymeric, protective matrices (Goswami et al., 2023). Testing EO activity solely in tryptic soy broth, on plastic surfaces, and against planktonic microbial cells fails to account for virtually all of these clinically relevant factors (Thaarup and Bjarnsholt, 2021).

Yet it is precisely within such complex wound environments that EOs may offer unique advantages - combining antibiofilm activity with immunomodulatory effects, and the capacity to penetrate tissue structures through both liquid diffusion and volatile dispersion.

To realize this potential, however, experimental *in vitro*/*ex vivo*/*in vivo* systems must be deliberately designed to reflect the wound milieu: not only in terms of medium composition, but also across all of the aforementioned experimental variables - including surface material, medium composition, incubation conditions, and spatial cell distribution.

Equally critical is the need to account for strain-level variability in microbial susceptibility to EOs. Even within a single species, individual isolates can differ dramatically in their biofilm architecture, metabolic activity, resistance phenotypes, and responsiveness to plant-derived compounds. Our studies and those of others have demonstrated that EO efficacy can span from near-complete eradication to functional resistance, depending on the strain tested. Also, the use of a single reference strain, however genetically tractable, cannot substitute for a representative panel encompassing clinical and environmental diversity. Moreover, a strain that appears susceptible to an EO under one set of laboratory parameters may exhibit tolerance under another (Ben Abdallah et al., 2020; Brożyna et al., 2021). No single test system can fully capture the complexity of real wound infections. Contrary, only a comparative approach, using multiple complementary models, can reveal patterns that would remain obscure in a reductionist setup and can provide translational outcomes.

Because integration of EOs into evidence-based antimicrobial practice remains a complex challenge, our group has already taken several key steps in this direction. In 2020, we introduced the AntiBioVol assay (Brożyna et al., 2020), which provided an experimental setup for testing volatile fractions in EO-mediated biofilm inhibition and exposed the interpretational flaws of classical aromatograms. In 2021, we demonstrated that *S. aureus* strains differ substantially in their susceptibility to EOs, emphasizing the need for broad strain panels (Brożyna et al., 2021). In 2022, we explored the biphasic (liquid and vapor) activity of EOs in structured, cellulose-based *Pseudomonas* biofilm models, underscoring how exposure format impacts antimicrobial efficacy (Brożyna et al., 2022). Most recently, we showed that the wound milieu itself - its fluid composition applied can dramatically alter EO performance, revealing context dependence at multiple biological levels (Brożyna et al., 2024). These studies are now complemented by the next work of ours (Brożyna et al., 2025), which examines the inter-and intraspecies variability of EO responses across diverse test systems.

Together, these efforts form the basis for a developing framework we refer to as Essential Oil Stewardship - an adaptable approach to studying and applying EOs in a scientifically grounded and translationally meaningful way. The present study serves as a practical demonstration of how EOs can be investigated using a structured, reproducible, and application-oriented setup, with the same methodological rigor expected of other anti-infective agents by examining strain-specific responses of *Staphylococcus aureus* to *Rosmarinus officinalis* L. and *Thymus vulgaris* L. EOs.

## 2 Materials and methods

### 2.1 Microorganisms

*Staphylococcus aureus* clinical (isolated from non-healing wounds) and reference strains were selected for the research. The strains were classified based on the analysis with MALDI-TOF (matrix-assisted laser desorption/ionization time-of-flight mass spectrometry) ultrafleXtreme spectrometer (Bruker, Billerica, USA) and identification with the Biotyper platform. All strains were previously deposited in the Strain and Line Collection of the Platform for Unique Models Application, Department of the Pharmaceutical Microbiology and Parasitology, Medical University of Wroclaw. The clinical ones were collected according to the bioethical approval of the following number: Bioethical Committee of Wroclaw Medical University, protocol #8/2016. In the first line of the experiment, for the biofilm mass and metabolic analysis, *S. aureus* ATCC 6538 (American Type Culture Collection) and twenty-five clinical strains were tested. The clinical strains included twelve MSSA (methicillin-susceptible *Staphylococcus aureus*) strains and thirteen MRSA (methicillin-resistant *Staphylococcus aureus*) isolates.

### 2.2 Essential oils

The antibiofilm activity of the following commercial essential oils was characterized:

- rosemary essential oil (REO), camphor chemotype, obtained from *Rosmarinus officinalis* L. leaves (Institute of Aromatherapy, Poland, batch number 1167/E4/74/T);
- thyme essential oil (TEO), thyme chemotype, obtained from the leaves of *Thymus vulgaris* L., Institute of Aromatherapy, Poland, batch number 33667/1/DW/46).

### 2.3 Culture conditions

Two different conditions were applied for biofilm culturing: medium and growth surface.

The used media included standard microbiological tryptic soy broth (TSB, Merck KGaA, Germany) and a medium reflecting the wound environment-in vitro wound milieu (IVWM).

According to the manufacturer’s specification, TSB is composed of (22092 Tryptic Soy Broth (TSB, (Tryptone Soya Broth, CASO Broth, Soybean Casein digest Broth, Casein Soya Broth), 2018):

- pancreatic digest of casein at a concentration of 17 g/L;
- dipotassium hydrogen phosphate at a concentration of 2.5 g/L;
- peptic digest of soybean at a concentration of 3 g/L;
- sodium chloride at a concentration of 5 g/L;
- glucose monohydrate at a concentration of 2.5 g/L.

Medium IVWM was prepared in compliance with the protocol developed by Kadam et al (Kadam et al., 2021):

Sterile, heat-inactivated fetal bovine serum (Biowest, France, cat. No. S181H) constituted 70% (v/v) of the medium volume. Other components were prepared as stock solutions, filtered with a 0.22 μm syringe filter or purchased sterile and added at the proper volume to reach the following final concentrations:

- 200–400 µg/mL fibrinogen from human plasma (Sigma-Aldrich, USA, cat. No. F3879; 10 mg/mL stock solution in 0.9% (w/v) sodium chloride (Stanlab, Poland));
- 20–30 µg/mL human lactoferrin (Sigma-Aldrich, USA, cat. No. L4040; 2 mg/mL stock solution in Dulbecco’s Phosphate Buffered Saline (Sigma-Aldrich, USA));
- 30–60 µg/mL human plasma fibronectin (Sigma-Aldrich, USA, cat. No. FC010; 1 mg/mL stock solution in autoclaved distilled water);
- 10–12 µg/mL collagen from bovine skin (concentration 2.9–3.2 mg/mL, Sigma-Aldrich, USA, cat. No. C4243);
- 11–12 mM lactic acid (Sigma-Aldrich, USA, cat. No. W261114; 11.4 M stock solution).

After combining all the components mentioned above, 0.9% (w/v) sodium chloride was added, constituting 19.5% (v/v) of the medium volume.

The medium was protected from light and stored at 2–8 °C for a maximum of seven days.

Two different surfaces were compared for biofilm growth: polystyrene (PS) or biocellulose (BC). In two performed tests (AntiBioVol (ABV, antibiofilm activity of volatile compounds method) and antibiofilm dressing’s activity measurement (A.D.A.M.)), biofilms were cultured on agar (A) instead of PS. 48-well polystyrene plates with the wells’ diameters of 11 mm (Wuxi Nest Biotechnology, China) served as the PS surface, and 2% (w/v) bacteriological lab agar (Biomaxima, Poland) was used to prepare agar discs (with a diameter of 11 mm). Biocellulose discs (with a diameter of 11 mm) were prepared in 48-well plates according to the following protocol.

*Komagataeibacter xylinus* ATCC 53524 strain was cultured in the Herstin–Schramm medium. The medium was composed of: 0.05% (w/v) MgSO4·7H_2_O (Chempur, Poland), 0.5% (w/v) bacto-peptone (VWR, USA), 2% (w/v) glucose (Chempur, Poland), 0.115% (w/v) citric acid monohydric (Chempur, Poland), 0.27% (w/v) Na_2_HPO_4_ (Stanlab, Poland), 0.5% (w/v) yeast extract (VWR, USA), and 1% (v/v) ethanol (Chempur, Poland). 50 µL of bacterial culture was added to the plates’ wells with 500 µL of the medium and incubated at 28°C for 7 days (under static conditions). After the formation of BC discs, they were removed from the wells and washed with a 0.1 M NaOH (Chempur, Poland) solution until all unadhered cells and cell debris were completely removed. Next, the BC discs were washed with double-distilled water until the pH neutralization. Sterilized in an autoclave, BC discs of the weight ranging from 0.15 g to 0.2 g were applied as the surface for biofilm growth.

### 2.4 Assessment of EOs’ chemical composition

The percentage composition of the EOs was evaluated using gas chromatography-mass spectrometry (GC–MS). Firstly, each oil was diluted 50 times in hexane, and then analysed with an Agilent 7890B GC system coupled with 7000GC/TQ, equipped with a PAL RSI85 autosampler (Agilent Technologies, USA) and an HP-5 MS column (30 m × 0.25 mm × 0.25 μm). Helium was applied as a carrier gas (total flow of 1 mL/min) and the ratio of injection splitting was 1:100. The initial temperature of was set at 50°C for 1 min, following with the temperature 170°C (4°C/min) and 280°C (10°C/min, kept for 2 min.). The MS detector settings were as follows: temperature of transfer line, source, and quadrupole – 320, 230, and 150°C, respectively, and 70 eV voltage of ionization. Detection was conducted in total scan mode at 30–400 m/z. The NIST 17.1 library and literature data were used to compare the obtained mass spectra and the retention index (RI). Indexes of linear retention were assessed under the conditions used for the EOs analysis with a mixture of C8–C20 saturated alkanes. The relative abundance of each component was presented as a percentage content based on peak area normalization (according to the MassHunter Workstation Software version B.09.00). The analysis was performed in three technical repetitions.

### 2.5 Biofilm culturing

For the experiments, staphylococcal biofilms were cultured according to the following procedure. Overnight bacterial cultures, incubated at 37°C in a specific medium (TSB or IVWM), were used for the preparation of suspensions for inoculation. A densitometer (DEN-1B, SIA Biosan, Latvia) was used to adjust the suspensions in saline to 0.5 MF (McFarland, 1.5 × 10^8^ CFU/mL (colony-forming unit)). Next, they were diluted to 1.5 × 10^5^ CFU/mL in the medium (TSB or IVWM) and added at a volume of 500 µL (unless otherwise stated) per well. For the test with PS as the growth surface, suspensions were poured directly into the wells of 48-well plates. For the two other surface types, the BC and A discs were previously soaked for 24 h in 500 µL of an appropriate medium. The inoculated plates were then incubated at 37 °C for 24 h under static conditions.

### 2.6 Biofilm features assessment

The staphylococcal biofilms were characterized by the assessment of the biomass (evaluated with crystal violet staining), metabolic activity (assessed with tetrazolium chloride staining), and the number of viable biofilm cells (evaluated with quantitative culturing). For the latter two tests, four variable conditions were applied: two types of media (TSB or IVWM) and two different surfaces (PS or BC). Due to methodological constraints, biofilm mass was assessed in both media but only on the polystyrene surface. The biofilms were cultured as described in the “Biofilm culturing” section.

#### 2.6.1 Biofilm mass evaluation

Once the biofilms were formed, the medium was removed, and 500 µL of 20% (v/v) crystal violet (Chempur, Poland) water solution was added for 10 min (room temperature). Next, the dye was gently pipetted out, and its excess was removed by one-step washing with 500 µL of saline. The plates were dried at 37°C for 10 min, and a 500 µL of 30% (v/v) acetic acid (Chempur, Poland) water solution was poured into the stained biofilm. The plates were shaken at 450 rpm (Mini-shaker PSU-2T, SIA Biosan, Latvia) for 30 min. at room temperature. Four samples, each at a volume of 100 µL, were taken from a separate testing well and transferred to the wells of 96-well plates (Wuxi Nest Biotechnology, China) for absorbance measurement at 550 nm with a spectrophotometer (MultiScan Go, Thermo Fisher Scientific, USA). The absorbance of the blank (medium incubated, stained, and extracted identically as the biofilms) was subtracted from the values of the tested samples. The mean of the four measurements was considered as a technical repetition. The experiment was performed in six technical repetitions and two biological repeats for each strain for a particular condition.

#### 2.6.2 Biofilm metabolic activity evaluation

Tetrazolium chloride (2,3,5-triphenyl-tetrazolium chloride, Sigma-Aldrich, USA) solution in the medium (TSB or IVWM) at the concentration of 0.1% (w/v) was applied as an indicator of metabolically-active cells and the produced formazan was extracted from the cells with mixture of methanol (Chempur, Poland) and acetic acid (9:1 ratio) (Chempur, Poland). For biofilms formed on the PS surface, the medium was removed from the biofilms, and the cells were stained with 500 µL of dye solution for 2 h/ 37°C. Subsequently, the dye was removed, and the plates were dried at 37°C for 10 min. The extractant was added at a volume of 500 µL directly to the wells with the biofilms, and the plates were subjected to shaking at 450 rpm for 30 min. at room temperature. For BC samples, the discs were transferred to fresh 48-well plates prior to the staining process and the extraction process was performed in 6-well plates (Wuxi Nest Biotechnology, China) using a 2 mL methanol:acetic acid solution. Four samples, each at a volume of 100 µL, were taken from a separate testing well (in PS and BC) and transferred to the wells of 96-well plates for the absorbance measurement at 490 nm using a spectrophotometer. The absorbance of the blank (medium incubated, stained, and extracted identically as the biofilms) was subtracted from the values of the tested samples. As the volume of the extracted solution differed between PS and BC samples, the PS results were adjusted by a dilution factor based on the calibration curve of the 1,3,5-triphenyltetrazolium formazan (Tokyo Chemical Industry, Japan) (data not presented). The mean of the four measurements was considered as a technical repetition. The experiment was performed in six technical repetitions and two biological repeats for each strain for a particular condition.

Based on the results of biofilm metabolic activity, ten *S. aureus* strains (five strains of high and five of low metabolic activity) were selected for further analysis of biofilm viable cell numbers and EOs antibiofilm activity.

#### 2.6.3 Number of biofilm viable cells

For the quantitative culturing test, 0.1% (w/v) saponin (VWR, USA) water solution was used. For the PS samples, the medium was first removed from the biofilms and the plates were shaken with 500 µL of the saponin solution for 30 s at 600 rpm. The solution was transferred to an Eppendorf tube (VWR, USA). Fresh 500 µL of saponin was added to the wells, and the shaking was repeated. The solution was then transferred to the Eppendorf tube. For biofilms formed on BC, the discs were transferred to 50 mL Falcon centrifuge tubes (Fl-Medical, Italy) and shaken for 1 min. with 10 mL of saponin. The decimal dilutions in saline were prepared from each sample (PS and BC) and cultured at a volume of 10 µL onto tryptic soy agar (Biomaxima, Poland) Petri dish plates (Noex, Poland). The plates were incubated for 24 h at 37°C. Single bacterial colonies were counted, and the number of viable cells was evaluated by calculating CFU/mL values. The experiment was performed in three technical repetitions and two biological repeats for each strain for a particular condition.

### 2.7 Antibiofilm activity of EOs evaluation

In the next part of the study, the antibiofilm activity of REO or TEO against staphylococcal biofilms was assessed using three complementary methods: the dilution method, AntiBioVol (ABV), and A.D.A.M. The ABV method evaluated EO activity in the volatile phase, while the dilution and A.D.A.M. methods assessed activity in the liquid phase. The biofilms were cultured as described in the section Biofilm culturing. The biofilms were incubated with the EOs for 24 h at 37°C under static conditions. The following controls were prepared: control of biofilm growth (C+, bacteria with medium or treated with saline), control of medium sterility (medium only), utility control (bacteria with the substance of proven antimicrobial activity). Separate plates were used for each EO and controls. In all methods, EOs activity was expressed as the percentage reduction of metabolically active biofilm cells after their staining with tetrazolium chloride. The biofilm cell reduction was calculated for each tested sample, considering mean growth control as 100% cell viability. All tests were performed in three technical repetitions and two biological repeats for each strain for a particular condition.

#### 2.7.1 Dilution method

In the experiment, the following three concentrations (v/v) of each EO were tested: 0.63%, 1.25%, and 2.5% of REO; 0.02%, 0.04%, and 0.08% of TEO. The maximum used EOs concentrations were selected due to complete or almost complete cell reduction at these concentrations in TSB/PS settings. The EOs were applied as emulsions in TSB or IVWM with Tween 20 (VWR, USA). The preliminary research demonstrated that the addition of Tween 20 at a concentration of 1% (v/v) did not influence staphylococcal growth (data not shown). Therefore, in the prepared stock emulsions, the emulsifier constituted 1% (v/v) of the emulsion volume. The concentration of REO and TEO in the stock emulsions was 2.5% (v/v) and 0.08% (v/v), respectively. In the first step of emulsion preparation, each EO was combined with Tween 20 and mixed with a magnetic stirrer at 1000 rpm (IKA RH basic 2, IKA, Germany) for 30 min. Next, the medium was added in four parts at five-minute intervals. Once the last part was added, the emulsion was stirred for 30 min. The consecutive EOs concentrations were prepared by the geometric dilution of each stock emulsion with the medium. The test was performed in 48-well plates, where the bacterial suspension was added directly to the wells (PS) or the wells with BC discs. After biofilm formation, the medium was removed from above the cells formed on PS and the BC discs were transferred to fresh 48-well plates. Next, EOs’ emulsions were added at a volume of 500 µL and the settings were incubated. The procedure of staining and absorbance measurement was carried out according to the protocol presented in the section Biofilm metabolic activity evaluation. The Octenisept (containing 0.1% of octenidine hydrochloride, and 2% of phenoxyethanol, Schülke & Mayr GmbH, Germany), diluted in the ratio 1:1 with TSB or IVWM, was used as the usability control. Bacteria in the medium were the growth control.

#### 2.7.2 AntiBioVol (antibiofilm activity of volatile compounds) method

The analysis was conducted according to the protocol presented in our previous study (Brożyna et al., 2020). For the research purpose, Petri dishes with 5 mm-thick agar and 24-well plates (Wuxi Nest Biotechnology, China) with the wells containing 1.5 mL of agar were prepared. Next, agar discs with a diameter of 11 mm were cut with a corkborer from agar Petri dishes and from the wells of the 24-well plate (the agar tunnels were then made in the wells). The early discs were kept and further used as agar (A) surface, the latter ones were discarded. Next, the agar discs and BC discs were placed in the wells of 24-well plates and soaked for 24 h with 500 µL of TSB or IVWM. The agar and BC discs were transferred to the agar tunnels in 24-well plates and placed at the bottom of them. 500 µL of the bacterial suspension was added to the wells, and the plates were incubated. Subsequently, the medium was removed from above the established biofilms. 500 µL of the tested EO was poured into the wells of fresh 24-well plates. Each biofilm-containing plate was placed upside down above the EO-containing plates. The plates were tightly connected with the parafilm and incubated for 24 h. Next, the plates were separated, and 400 µL of 0.1% (w/v) tetrazolium chloride solution was added for 2 h (37°C). Next, the plates were opened and placed in the incubator with a fan at 80°C until the solution above the biofilm evaporated. The discs were transferred to 6-well plates, and formazan extraction was performed with 4 mL of the extracting solution. Three samples at the volume of 100 µL were taken from a separate testing well (in A and BC) and transferred to the wells of 96-well plates for the absorbance measurement at 490 nm with a spectrophotometer. The absorbance of the blank (medium incubated, stained and extracted identically as the biofilms) was subtracted from the values of the tested samples. 0.9% (w/v) sodium chloride was used as the growth control; 96% (v/v) ethanol (Chempur, Poland) was used as usability control.

#### 2.7.3 A.D.A.M. (antibiofilm dressing’s activity measurement) method

The test was performed following the procedure described in our previous paper (Junka et al., 2017). Biocellulose dressings were applied as the carriers for EOs. Biocellulose dressings were prepared using the methodology described in the section “Culture conditions” for BC disc preparation. However, for the biocellulose dressings, 24-well plates with 2 mL of the Herstin–Schramm and inoculated with 200 µL of *K. xylinus* were used. The biocellulose dressings of the weight range from 0.55 g to 0.65 g were selected. Twenty-four hours before the application, the dressings were placed in the 24-well plates and soaked with 1 mL of the EO. Therefore, the concentration of EO after the soaking was calculated as follows:

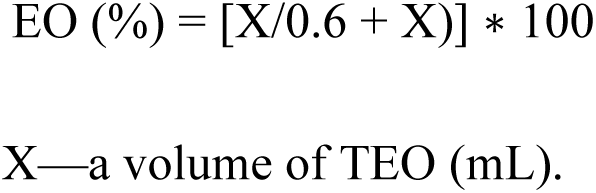

The plates were sealed with parafilm and stored at 2-8°C. For research purposes, biofilms were cultured in TSB or IVWM and on A or BC surfaces. The process of surface preparation and biofilm cultivation was presented in the section AntiBioVol (antibiofilm activity of volatile compounds) method. After the biofilm’s development, the medium was gently removed from above the cells and replaced with 600 µL of fresh medium. The biocellulose dressings were then placed on top of the wells, and the plates were incubated for 24 h. Next, the biocellulose dressings were discarded, 200 µL of the medium above the biofilm was removed, and 100 µL of 0.5% (w/v) tetrazolium chloride was added for 2 h (37°C). The following steps of the AniBioVol methodology were performed in the same manner. 0.9% (w/v) sodium chloride was used as the growth control; Prontosan Wound Irrigation Solution (composed of 0.1% polyhexamethylene biguanide and 0.1% undecylenamidopropyl betaine, B. Braun Melsungen AG, Germany), was used as usability control.

### 2.8 Fluorescence microscopy

To illustrate the differences in the antibiofilm effects of rosemary (REO) and thyme (TEO) essential oils under varying nutritional conditions, representative MSSA strains S3 and S4 were selected for fluorescence-based imaging. The strains were selected based on the AntiBioVol results obtained for TSB medium. S4 was the least susceptible to REO and TEO on both A and PS surfaces. S3 was among the most susceptible strains under each condition and EOs. The biofilms were cultured on PS in two media - TSB or IVWM - and treated with REO or TEO following the protocols described in the AntiBioVol section, with minor modifications to accommodate microscopic analysis. The experiment was conducted using 24-well plates (Wuxi Nest Biotechnology, China), with 1 mL of bacterial suspension applied per well (wells without agar tunnels). Following incubation with essential oils or saline (growth control, C+), the biofilms were stained using the Filmtracer™ LIVE/DEAD™ Biofilm Viability Kit (Thermo Fisher Scientific, USA), prepared according to the manufacturer’s instructions. A volume of 10 μL of the staining reagent was added to each well and incubated at room temperature in the dark for 15 minutes. Subsequently, the wells were gently rinsed with 200 μL of double-distilled water and fixed with 4% (v/v) formaldehyde (Chempur, Poland) for 2 h. The analysis was performed using a fluorescence microscope, Etaluma 600 LumaScope (object lens with 4× magnification, Etaluma, USA).

### 2.9 Analysis of biofilms’ protein profiles

The changes of bacterial proteins after the exposition to EOs’ volatile fractions were detected using mass spectrometry measurement on a MALDI-TOF (matrix-assisted laser desorption/ionization time-of-flight mass spectrometry) ultrafleXtreme spectrometer (Bruker, USA) and identification with the Biotyper platform. This procedure identifies microorganisms by comparing the mass spectra of microbial proteins, extracted from bacterial colonies, with an extensive library of 8,468 reference spectra. Biofilm culturing and treatment with EOs were performed in TSB or IVWM medium and on BC surface, following the protocols described in the AntiBioVol section with S3 and S4 strains. Subsequently, BC disks with biofilms treated with EOs or growth control samples (C+) were ground before the actual extraction. BC were ground using a microtube homogenizer (D1030-E, BeadBug, Benchmark Scientific, Inc., USA) in a dedicated tube with 400 µL water and 1 mm diameter glass beads for 2 min. at a shaking intensity of 4000 rpm. Then, the ground BC was centrifuged for 30 s at 4000 rpm and 4°C (Centrifuge 5424R, Eppendorf, VWR, USA). 300 µL of supernatant with bacterial suspension was further extracted according to the standard procedure (Dudek et al., 2019). Next, the samples were subjected to extraction using a mixture of formic acid/acetonitrile (Chempur, Poland). After proteins extraction 1 µl of sample was spotted on MALDI target plate, dried and covered with 1 µl of matrix solution (10 mg/mL HCCA (α-cyano-4-hydroxycinnamic acid, VWR, USA) in acetonitrile:water:trifluoroacetic acid solution (50:47.5:2.5, v:v:v) (Chempur, Poland). The investigated bacterial strains were reliably identified as *Staphylococcus aureus*, with a score value of ≥2.000 serving as the threshold for species identification. The protein profiles presented as mass spectra were generated from a single technical repetition.

### 2.10 Statistical analysis

All statistical analyses and visualizations were conducted using R (version 4.4.3) via RStudio (2025-02-13), utilizing packages including ggplot2, dplyr, rstatix, and Dunn’s test. Data preprocessing included outlier removal based on the standard thresholding method at 1.5 × the interquartile range (IQR). Normality of data distribution and homogeneity of variance were assessed both visually and using the Shapiro–Wilk and Levene’s tests, respectively.

To evaluate differences in biofilm activity of *Staphylococcus aureus* strains across different media and surfaces, the Kruskal–Wallis test was employed, followed by post-hoc pairwise comparisons using Dunn’s test with Bonferroni correction. The Bonferroni method was chosen to reduce the risk of false positives that can arise when conducting many related statistical tests. The same statistical approach was applied to assess the antibiofilm activity of essential oils (EOs) against *S. aureus* strains. Statistical significance was considered at *p* < 0.05.

Correlations between biofilm biomass, metabolic activity, and viable cell number (CFU/mL) were assessed using correlation plots with linear regression lines of best fit.

## 3 Results

### 3.1 Evaluation of EOs’ chemical composition

In the initial phase of the investigation, the percentage contents of specific compounds in the EOs were evaluated using GC-MS methodology (Supplementary Table S1) and compared to their ranges specified in the European Pharmacopoeia XI standards. Twenty-nine compounds were found in REO; four of them constituted 68.38% of this EO content, α-pinene (21.07%), 1,8-cineole (19.98%), camphor (18.52%), and camphene (8.81%). The p-cymene content was not aligned with the European Pharmacopoeia XI standards; however, it exceeded the indicated level by just 0.26%. The TEO’s most abundant components were thymol, p-cymene, g-terpinene, and carvacrol, constituting 50.59%, 19.2%, 9.06%, and 5.56% of this EO content, respectively. Only two compounds deviated from the pharmacopeial standards: carvacrol slightly exceeded the upper limit by 0.15%, while carvacrol methyl ether was not detected.

### 3.2 Comparison of biofilm features

The ability of *S. aureus* strains to form biofilms under the tested conditions was evaluated by analyzing three parameters: biofilm mass, metabolic activity, and viable cell number. Biofilms were cultured in two media-TSB and IVWM - and, for the assessment of metabolic activity and viable cell number, also on two types of surfaces: polystyrene and bacterial cellulose. Accordingly, these biofilm parameters were compared under four distinct growth conditions when both variables (medium and surface) were considered, or grouped into two conditions when only one variable (either medium or surface) was compared. Due to methodological constraints, biofilm mass was assessed in both media but only on the polystyrene surface; for this parameter, the comparison was limited to the two media. All tested strains formed biofilms under the applied conditions (Figures 1–3, Tables 1– and 2, Supplementary Figures S1–S3, Supplementary Tables S2–S7). The distributions of each parameter are presented in Supplementary Figures S1–S3 and Supplementary Tables S2, S4, and S6.

**Figure 1.**
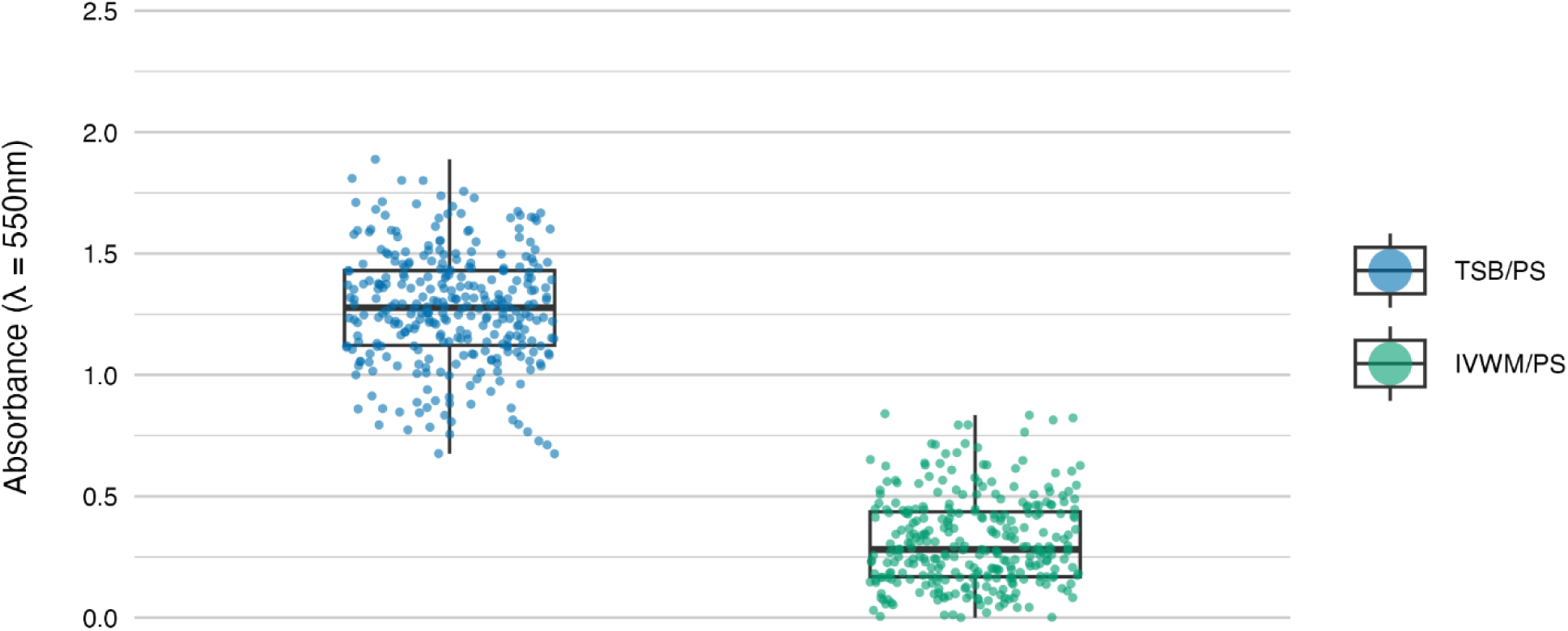
Biofilm mass of *S. aureus* strains (n=26) cultured on polystyrene (PS) surface and in different media: tryptic soy broth (TSB) or in in vitro wound milieu (IVWM). Each box displays the interquartile range (IQR; 25th to 75th percentiles), with the bold horizontal line indicating the median. Whiskers extend to the most extreme data points within 1.5 × IQR from the lower and upper quartiles. Technical repetitions are shown as dots. Differences were statistically significant, p ≤ 0.0001. Dunn’s test was performed. Adjusted p value includes Bonferroni correction (Supplementary Table S3).

The mean level of biofilm mass formed in TSB was four times higher than in IVWM, with a level of statistical significance of p ≤ 0.0001 (Dunn’s test. Adjusted p value includes Bonferroni correction, Figure 1, Supplementary Tables S2 and S3).

Biofilm metabolic activity was significantly different between the four growing conditions (p ≤ 0.0001, Dunn’s test. Adjusted p value includes Bonferroni correction. Figure 2, Table 1, Supplementary Tables S4 and S5). The highest difference was observed between TSB/PS and IVWM/PS (7.5 times higher in TSB/PS). The mean metabolic activity in IVWM/BC was 3 times higher than in IVWM/PS, and 1.6 times higher in TSB/PS than in TSB/BC. When comparing surface type regardless of medium, mean metabolic activity was similar for PS and BC (1.8 vs. 1.6, respectively). In contrast, when comparing the media regardless of surface, biofilms cultured in TSB showed approximately threefold higher metabolic activity than those in IVWM, with strong statistical significance (p ≤ 0.0001, Dunn’s test. Adjusted p value includes Bonferroni correction).

**Figure 2.**
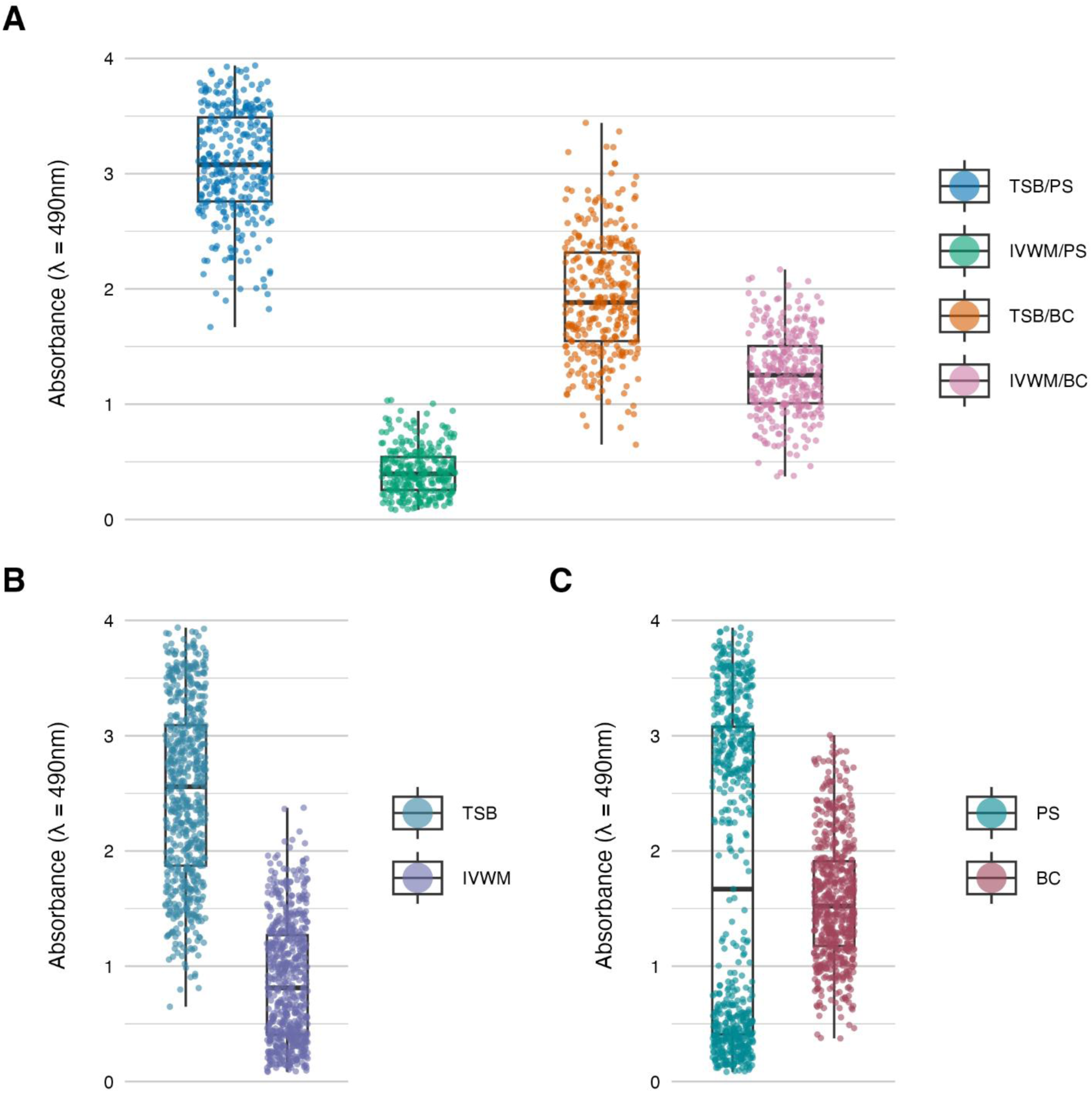
Biofilm metabolic activity of *S. aureus* strains (n=26) cultured on different surfaces: polystyrene (PS) or biocellulose (BC), and in different media: tryptic soy broth (TSB) or in in vitro wound milieu (IVWM). **(A)** Dividing condition: medium and surface. **(B)** Dividing condition: medium. **(C)** Dividing condition: surface. Each box displays the interquartile range (IQR; 25th to 75th percentiles), with the bold horizontal line indicating the median. Whiskers extend to the most extreme data points within 1.5 × IQR from the lower and upper quartiles. Technical repetitions are shown as dots.

**Figure 3.**
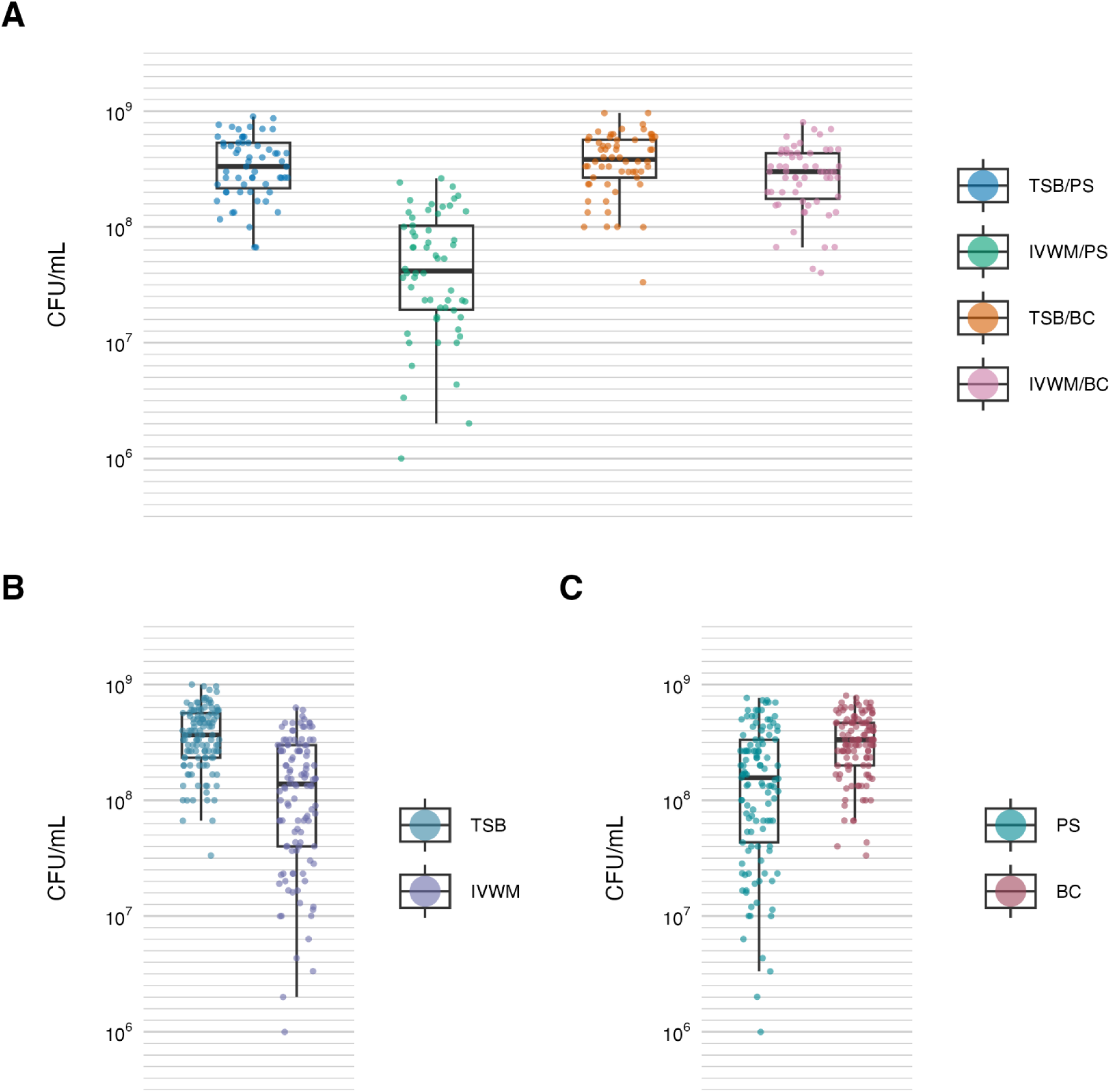
Biofilm viable cell number of *S. aureus* strains (n=10) cultured on different surfaces: polystyrene (PS) or biocellulose (BC), and in different media: tryptic soy broth (TSB) or in in vitro wound milieu (IVWM). **(A)** Dividing condition: medium and surface. **(B)** Dividing condition: medium. **(C)** Dividing condition: surface. Each box displays the interquartile range (IQR; 25th to 75th percentiles), with the bold horizontal line indicating the median. Whiskers extend to the most extreme data points within 1.5 × IQR from the lower and upper quartiles. Technical repetitions are shown as dots. CFU/mL-colony-forming unit/mL.

**Table 1.** Summarized statistical differences in biofilm metabolic activity of *S. aureus* strains (n=26) cultured on different surfaces: polystyrene (PS) or biocellulose (BC), and in different media: tryptic soy broth (TSB) or in in vitro wound milieu (IVWM). **(A)** Dividing condition: medium and surface. (B) Dividing condition: medium or surface. Dunn’s test was performed. Adjusted p value includes Bonferroni correction. Values of p<0.05 were considered significant, p ≤ 0.0001 was marked with four asterisks. Ns-no significant differences. X-not applicable.

|  | TSB/PS | IVWM/PS | TSB/BC | IVWM/BC |
| --- | --- | --- | --- | --- |
| TSB/PS | X | **** | **** | **** |
| IVWM/PS | **** | X | **** | **** |
| TSB/BC | **** | **** | X | **** |
| IVWM/BC | **** | **** | **** | X |

**(B)**
|  | IVWM | BC |
| --- | --- | --- |
| TSB | **** | X |
| PS | X | ns |

Similarly to metabolic activity, the mean number of viable biofilm cells was lowest under the IVWM/PS condition. Notably, this was the only condition that differed significantly from all three remaining growth setups (p ≤ 0.0001, Dunn’s test. Adjusted p value includes Bonferroni correction). When grouped by individual variables, biofilms cultured in TSB contained significantly more viable cells than those grown in IVWM (p ≤ 0.0001, Dunn’s test. Adjusted p value includes Bonferroni correction), and biofilms formed on BC surfaces showed significantly higher viable cell numbers than those formed on PS (p ≤ 0.0001, Dunn’s test. Adjusted p value includes Bonferroni correction, Figure 3, Table 2, Supplementary Tables S6 and S7).

**Table 2.** Summarized statistical differences in biofilm viable cell number of *S. aureus* strains (n=10) cultured on different surfaces: polystyrene (PS) or biocellulose (BC), and in different media: tryptic soy broth (TSB) or in in vitro wound milieu (IVWM). **(A)** Dividing condition: medium and surface. **(B)** Dividing condition: medium or surface. Dunn’s test was performed. Adjusted p value includes Bonferroni correction. Values of p<0.05 were considered significant, p ≤ 0.0001 was marked with four asterisks. Ns-no significant differences. X-not applicable.

|  | **TSB/PS** | **IVWM/PS** | **TSB/BC** | **IVWM/BC** |
| --- | --- | --- | --- | --- |
| **TSB/PS** | **X** | **\*\*\*\*** | **ns** | **ns** |
| **IVWM/PS** | **\*\*\*\*** | **X** | **\*\*\*\*** | **\*\*\*\*** |
| **TSB/BC** | **ns** | **\*\*\*\*** | **X** | **ns** |
| **IVWM/BC** | **ns** | **\*\*\*\*** | **ns** | **X** |

(B)
|  | **IVWM** | **BC** |
| --- | --- | --- |
| **TSB** | **\*\*\*\*** | **X** |
| **PS** | **X** | **\*\*\*\*** |

In the next step of the study, we have analyzed the correlation between the features of biofilms cultured under the tested conditions (Figures 4-6). The highest and positive linear correlation was observed for biofilm mass and metabolic activity in IVWM/PS (R^2^=0.67, Figure 4A). However, regardless of the dividing condition, no strong correlation was demonstrated between the biofilm characteristics.

**Figure 4.**
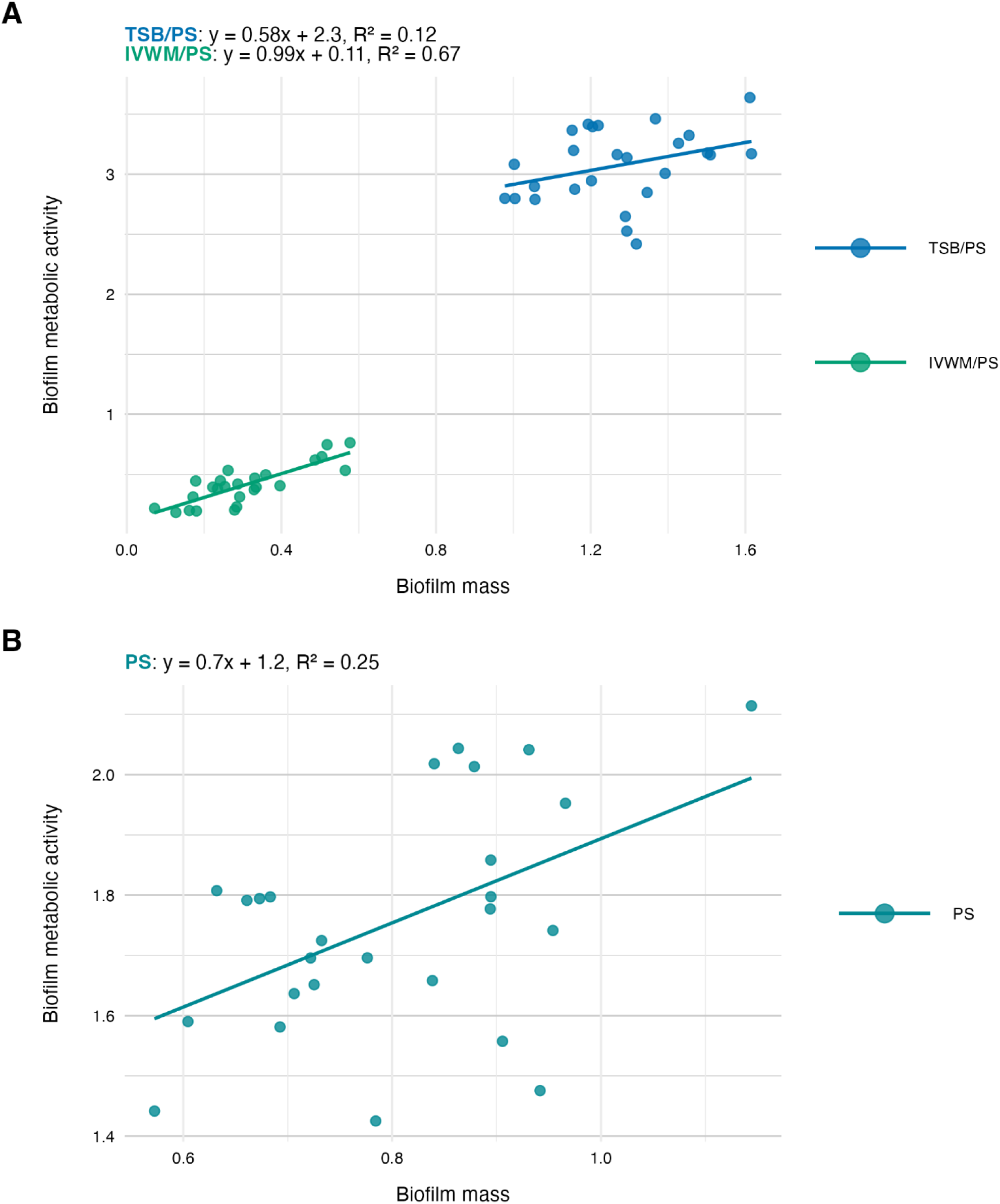
Scatter plots of correlations of *S. aureus* strains’ (n=26) biofilm mass and biofilm metabolic activity. Biofilms were cultured on polystyrene (PS) and in different media: tryptic soy broth (TSB) or in in vitro wound milieu (IVWM). **(A)** Dividing condition: medium. **(B)** Dividing condition: surface. The points denote the means for each strain. Data fitted on a linear trend line. The equation for the line of best fit and R^2^-coefficient of determination are presented in the top left corners of the respective panels.

**Figure 5.**
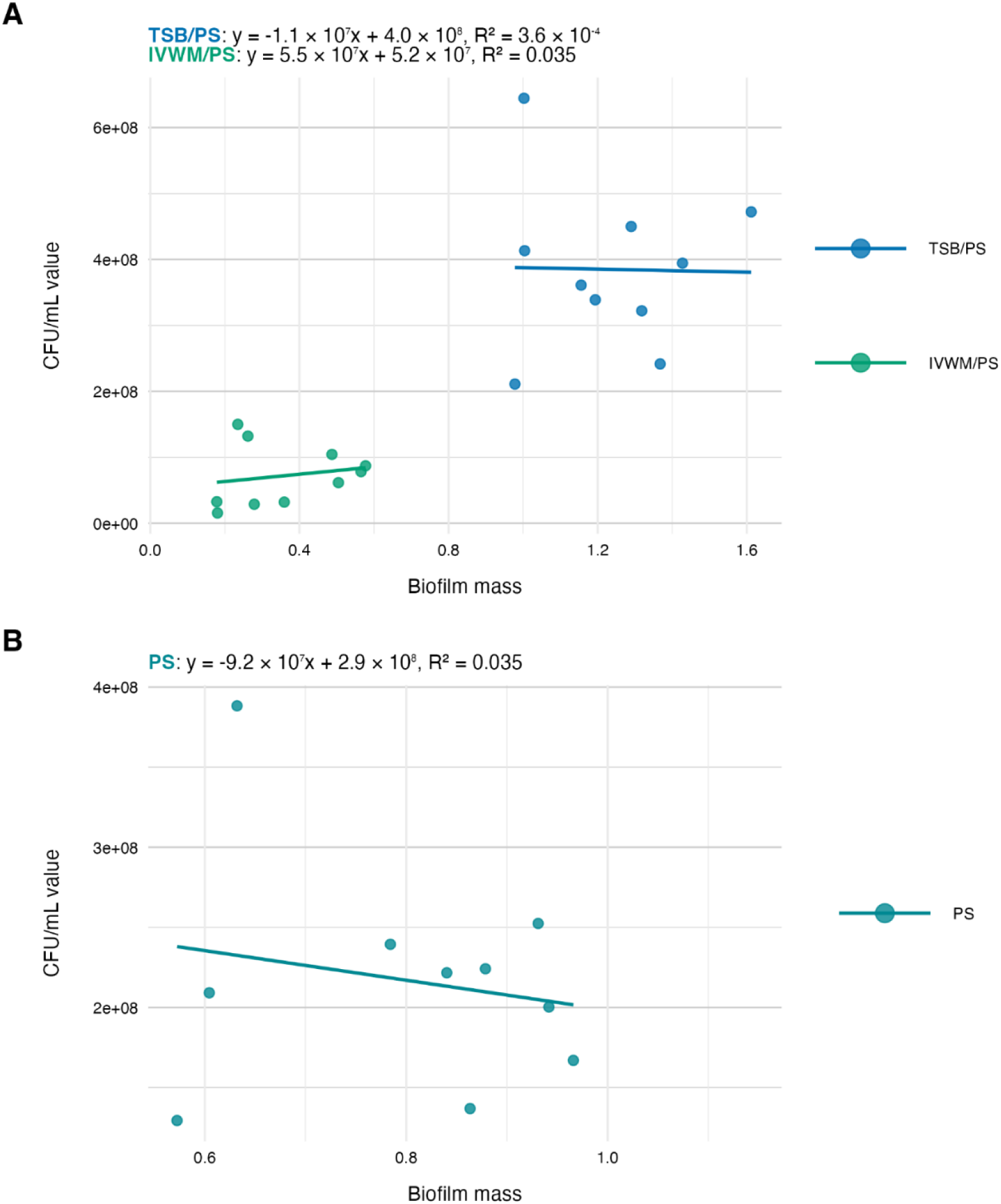
Scatter plots of correlations of *S. aureus* strains’ (n=10) biofilm mass and biofilm viable cell number (CFU/mL-colony-forming unit/mL). Biofilms were cultured on polystyrene (PS) and in different media: tryptic soy broth (TSB) or in in vitro wound milieu (IVWM). **(A)** Dividing condition: medium. **(B)** Dividing condition: surface. The points denote the means for each strain. Data fitted on a linear trend line. The equation for the line of best fit and R^2^-coefficient of determination are presented in the top left corners of the respective panels.

### 3.3 Comparison of antibiofilm activity of the EOs

Subsequently, the antibiofilm activity of rosemary (REO) and thyme (TEO) EOs was assessed using three complementary methods: the dilution method, AntiBioVol (ABV), and A.D.A.M. The ABV method evaluated EO activity in the volatile phase, while the dilution and A.D.A.M. methods assessed activity in the liquid phase (Figures 7 and 8, Tables 3-7, Supplementary Figures S4-S6 and Supplementary Tables S8-S19). All tests were performed under four different growth conditions, combining two media (TSB and IVWM) with two surface types - polystyrene (for the dilution method) or agar (for ABV and A.D.A.M.), and bacterial cellulose. Accordingly, comparisons were made across four distinct combinations when both variables (medium and surface) were considered, or grouped by a single variable (medium or surface) when assessing isolated effects.

**Figure 6.**
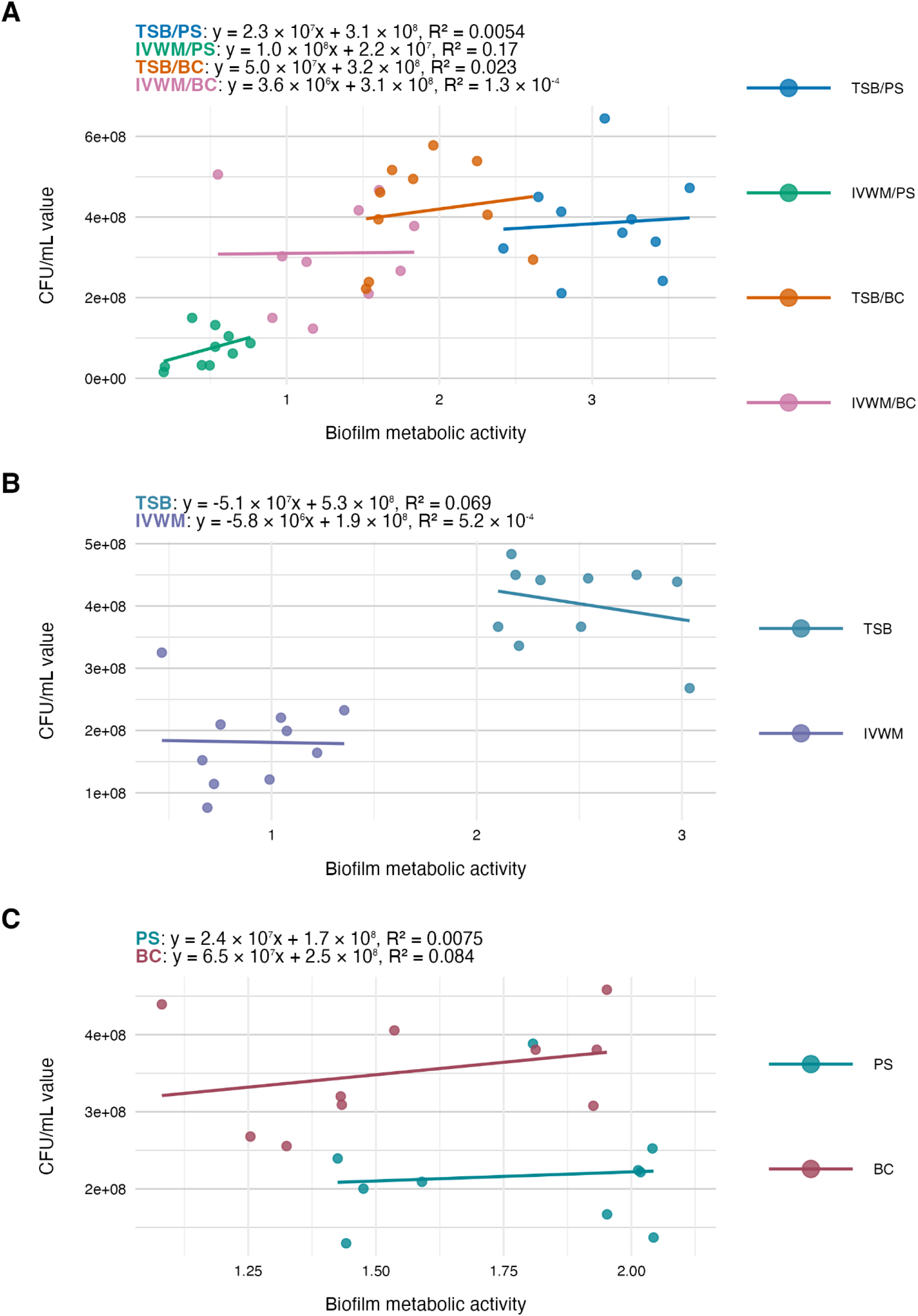
Scatter plots of correlations of *S. aureus* strains’ (n=10) biofilm metabolic activity and biofilm viable cell number (CFU/mL-colony-forming unit/mL). Biofilms were cultured on different surfaces: polystyrene (PS) or biocellulose (BC), and in different media: tryptic soy broth (TSB) or in in vitro wound milieu (IVWM). **(A)** Dividing condition: medium and surface. **(B)** Dividing condition: medium. **(C)** Dividing condition: surface. The points denote the means for each strain. Data fitted on a linear trend line. The equation for the line of best fit and R^2^-coefficient of determination are presented in the top left corners of the respective panels.

**Figure 7.**
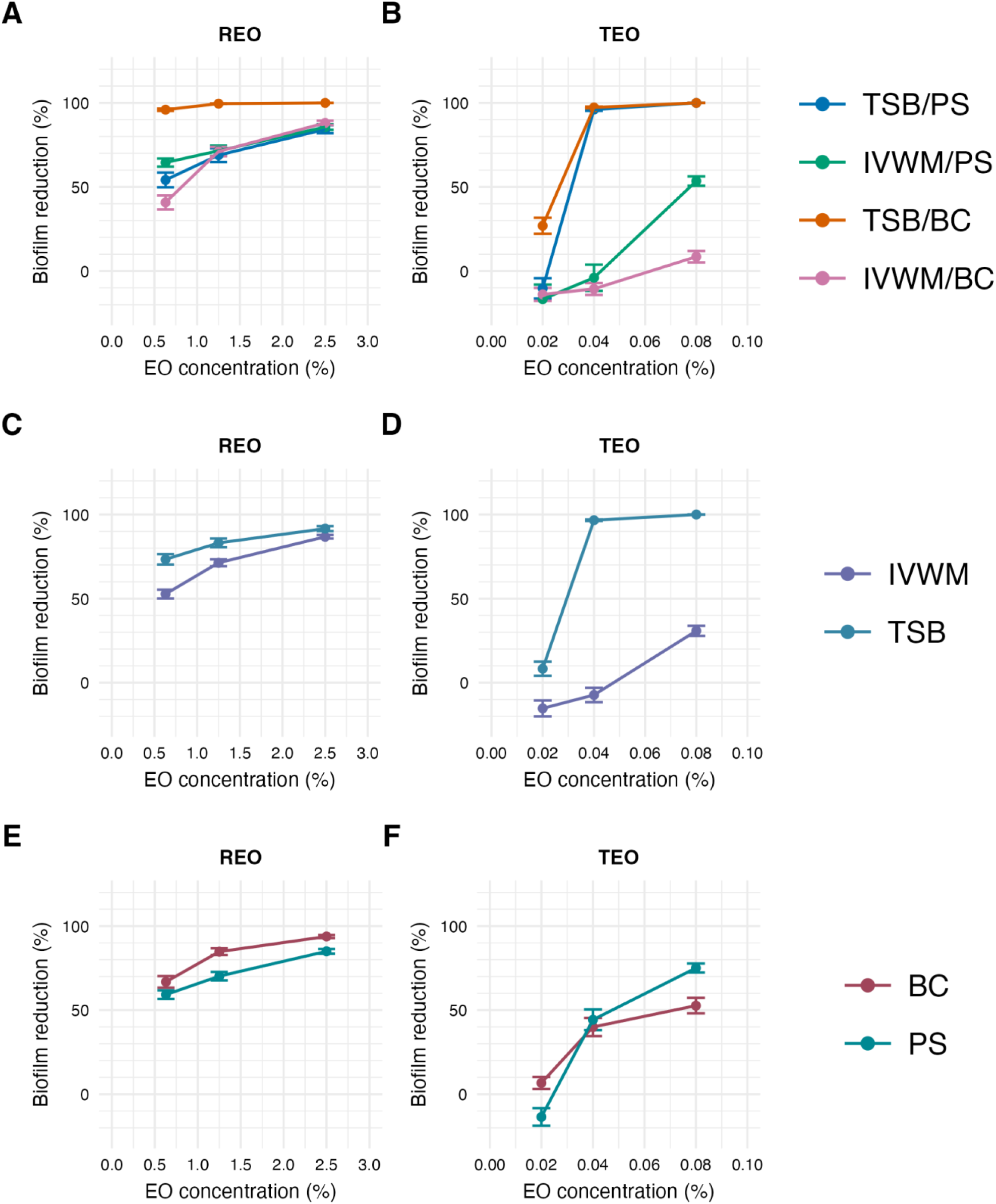
Antibiofilm activity of rosemary essential oil (REO) or thyme essential oil (TEO) against *S. aureus* strains (n=10) cultured on different surfaces: polystyrene (PS) or biocellulose (BC) and in different media: tryptic soy broth (TSB) or in in vitro wound milieu (IVWM), assessed with a dilution method. **(A, B)** Dividing condition: medium and surface. **(C, D)** Dividing condition: medium. **(E, F)** Dividing condition: surface. The points denote the mean value, and the error lines denote the standard error of measurement.

**Figure 8.**
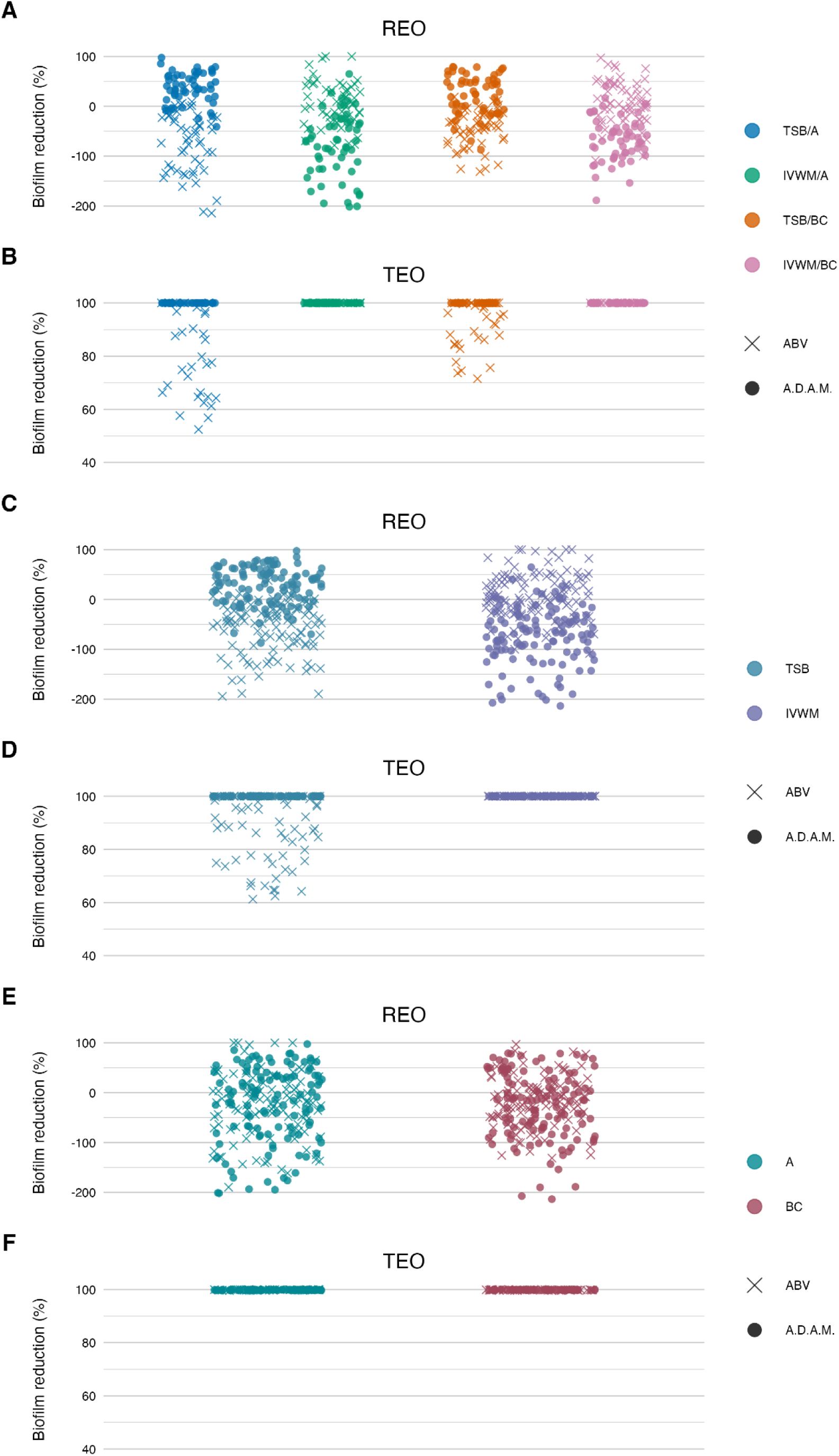
Antibiofilm activity of rosemary essential oil (REO) or thyme essential oil (TEO) against *S. aureus* strains (n=10) cultured on different surfaces: agar (A) or biocellulose (BC) and in different media: tryptic soy broth (TSB) or in in vitro wound milieu (IVWM), assessed with an antibiofilm activity of volatile compounds (ABV) or an antibiofilm dressing’s activity measurement (A.D.AM.) methods. **(A, B)** Dividing condition: medium and surface. **(C, D)** Dividing condition: medium. **(E, F)** Dividing condition: surface. The points denote technical repetitions.

**Table 3.** Summarized statistical differences in antibiofilm activity of rosemary essential oil (REO) against *S. aureus* strains (n=10) cultured on different surfaces: polystyrene (PS) or biocellulose (BC) and in different media: tryptic soy broth (TSB) or in in vitro wound milieu (IVWM), assessed with a dilution method. **(A, B, C)** Dividing condition: medium and surface. **(D)** Dividing condition medium or surface; results were compared between the same EO’s concentrations. Dunn’s test was performed. Adjusted p value includes Bonferroni correction. Values of p<0.05 were considered significant, p ≤ 0.05 was marked with one asterisk, p ≤ 0.01 was marked with two asterisks, p ≤ 0.0001 was marked with four asterisks. Ns-no significant differences. X-not applicable.

|  | 0.63% REO |  |  |  |
| --- | --- | --- | --- | --- |
|  | TSB/PS | IVWM/PS | TSB/BC | IVWM/BC |
| TSB/PS | X | ns | **** | ns |
| IVWM/PS | ns | X | **** | * |
| TSB/BC | **** | **** | X | **** |
| IVWM/BC | ns | * | **** | X |

(B)
|  | 1.25% REO |  |  |  |
| --- | --- | --- | --- | --- |
|  | TSB/PS | IVWM/PS | TSB/BC | IVWM/BC |
| TSB/PS | X | ns | **** | ns |
| IVWM/PS | ns | X | **** | ns |
| TSB/BC | **** | **** | X | **** |
| IVWM/BC | ns | ns | **** | X |

(C)
|  | **2.5% REO** |  |  |  |
| --- | --- | --- | --- | --- |
|  | **TSB/PS** | **IVWM/PS** | **TSB/BC** | **IVWM/BC** |
| **TSB/PS** | **X** | **ns** | **\*\*\*\*** | **ns** |
| **IVWM/PS** | **ns** | **X** | **\*\*\*\*** | **ns** |
| **TSB/BC** | **\*\*\*\*** | **\*\*\*\*** | **X** | **\*\*\*\*** |
| **IVWM/BC** | **ns** | **ns** | **\*\*\*\*** | **X** |

(D)
| **REO** |  |  |  |
| --- | --- | --- | --- |
| **EO concentration** |  | **IVWM** | **BC** |
| **0.63%** | **TSB** | **\*\*\*\*** | **X** |
| **0.63%** | **PS** | **X** | **\*\*** |
| **1.25%** | **TSB** | **\*\*\*\*** | **X** |
| **1.25%** | **PS** | **X** | **\*\*\*\*** |
| **2.5%** | **TSB** | **\*\*\*\*** | **X** |
| **2.5%** | **PS** | **X** | **\*\*\*\*** |

According to all of the performed tests (Figures 7 and 8, Supplementary Tables S8, S11-S12, S15-S16, S18), TEO exhibited higher antimicrobial activity than REO under all applied conditions. Complete biofilm eradication was assessed after the treatment with octenidine hydrochloride and ethanol (usability controls in dilution and ABV methods, respectively) under all growth conditions. The following values of biofilm reduction were observed after the exposure to polyhexanide (usability control in the A.D.A.M. method): 27% in TSB/A, 17% in TSB/BC, and 19% in IVWM/A and IVWM/BC settings.

In the dilution method, REO in TSB/BC setting was the most effective against *S. aureus* biofilm at each of the tested concentrations, and the activity was significantly higher than the three other conditions (p ≤ 0.0001, Dunn’s test. Adjusted p value includes Bonferroni correction. Figure 7, Table 3, Supplementary Tables S8 and S9). The highest differences in mean biofilm reduction between conditions at the same REO concentration were 55% (TSB/BC vs IVWM/BC at 0.63% (v/v)). Antibiofilm activity of REO was also significantly higher in TSB than in IVWM (p ≤ 0.0001, Dunn’s test. Adjusted p value includes Bonferroni correction) and on BC than PS (p ≤ 0.01 at concentration of 0.63% (v/v); p ≤ 0.0001 at concentrations of 1.25% (v/v) and 2.5% (v/v), Dunn’s test. Adjusted p value includes Bonferroni correction), regardless of the concentration.

Analysis of TEO antibiofilm activity under two dividing conditions revealed the highest biofilm reduction in TSB/BC at all concentrations (similar to REO) and the lowest in IVWM/BC at concentrations of 0.04% (v/v) and 0.08% (v/v) (Figure 7, Table 4, Supplementary Tables S8 and S10). Although in these settings not all differences were statistically significant, the mean biofilm eradication > 96% was assessed at the oil concentration of 0.08% (v/v) and 0.04% (v/v) in TSB/BC and TSB/PS, while in IVWM/BC the mean reduction was 9% and-11% (indicating higher viability of cells treated with the EO than in the growth control) for 0.08% (v/v) and 0.04% (v/v), respectively. TEO exhibited significantly higher activity in TSB than in IVWM at all tested concentrations (p ≤ 0.0001 at concentrations of 0.08% (v/v) and 0.04% (v/v); p ≤ 0.001 at concentration of 0.02%, Dunn’s test. Adjusted p value includes Bonferroni correction).

**Table 4.** Summarized statistical differences in antibiofilm activity of thyme essential oil (TEO) against *S. aureus* strains (n=10) cultured on different surfaces: polystyrene (PS) or biocellulose (BC) and in different media: tryptic soy broth (TSB) or in in vitro wound milieu (IVWM), assessed with a dilution method. **(A, B, C)** Dividing condition: medium and surface. **(D)** Dividing condition medium or surface; results were compared between the same EO’s concentrations. Dunn’s test was performed. Adjusted p value includes Bonferroni correction. Values of p<0.05 were considered significant, p ≤ 0.05 was marked with one asterisk, p ≤ 0.01 was marked with two asterisks, p ≤ 0.001 was marked with three asterisks, p ≤ 0.0001 was marked with four asterisks. Ns-no significant differences. X-not applicable.

|  | 0.02% TEO |  |  |  |
| --- | --- | --- | --- | --- |
|  | TSB/PS | IVWM/PS | TSB/BC | IVWM/BC |
| TSB/PS | X | ns | **** | ns |
| IVWM/PS | ns | X | *** | ns |
| TSB/BC | **** | *** | X | **** |
| IVWM/BC | ns | ns | **** | X |

(B)
|  | 0.04% TEO |  |  |  |
| --- | --- | --- | --- | --- |
|  | TSB/PS | IVWM/PS | TSB/BC | IVWM/BC |
| TSB/PS | X | **** | ns | **** |
| IVWM/PS | **** | X | **** | ns |
| TSB/BC | ns | **** | X | **** |
| IVWM/BC | **** | ns | **** | X |

(C)
|  | 0.08% TEO |  |  |  |
| --- | --- | --- | --- | --- |
|  | TSB/PS | IVWM/PS | TSB/BC | IVWM/BC |
| TSB/PS | X | **** | ns | **** |
| IVWM/PS | **** | X | **** | *** |
| TSB/BC | ns | **** | X | **** |
| IVWM/BC | **** | *** | **** | X |

|  | TEO |  |  |
| --- | --- | --- | --- |
| EO concentration |  | IVWM | BC |
| 0.02% | TSB | *** | X |
| 0.02% | PS | X | * |
| 0.04% | TSB | **** | X |
| 0.04% | PS | X | ns |
| 0.08% | TSB | **** | X |
| 0.08% | PS | X | ** |

According to the ABV, volatile forms of REO displayed low antibiofilm activity, or the metabolic activity of cells treated with the oil was higher than in growth control in all applied settings. Significantly higher mean reduction was observed in IVWM (10%) than in TSB (-58%) (p ≤ 0.0001, Dunn’s test. Adjusted p value includes Bonferroni correction), with simultaneously no significant differences between the A and BC surfaces (Figure 8, Table 5, Supplementary Tables S12 and S13). In all four conditions, TEO reduced staphylococcal biofilm in at least 88% (TSB/A). The reduction was also significantly higher in IVWM than in TSB (p ≤ 0.0001, Dunn’s test. Adjusted p value includes Bonferroni correction); however, the equal level of biofilm reduction was demonstrated regarding the surface (A or BC) (Figure 8, Table 5, Supplementary Tables S12 and S14).

**Table 5.** Summarized statistical differences in antibiofilm activity of rosemary essential oil (REO) or thyme essential oil (TEO) against *S. aureus* strains (n=10) cultured on different surfaces: agar (A) or biocellulose (BC) and in different media: tryptic soy broth (TSB) or in in vitro wound milieu (IVWM), assessed with an antibiofilm activity of volatile compounds method. **(A)** Dividing condition: medium and surface. **(B)** Dividing condition: medium or surface. Dunn’s test was performed. Adjusted p value includes Bonferroni correction. Values of p<0.05 were considered significant, p ≤ 0.0001 was marked with four asterisks. Ns-no significant differences. X-not applicable.

|  | REO |  |  |  | TEO |  |  |  |
| --- | --- | --- | --- | --- | --- | --- | --- | --- |
|  | TSB/A | IVWM/A | TSB/BC | IVWM/BC | TSB/A | IVWM/A | TSB/BC | IVWM/BC |
| TSB/A | X | **** | ns | **** | X | **** | ns | **** |
| IVWM/A | **** | X | **** | ns | **** | X | **** | ns |
| TSB/BC | ns | **** | X | **** | ns | **** | X | **** |
| IVWM/BC | **** | ns | **** | X | **** | ns | **** | X |

(B)
|  | REO |  | TEO |  |
| --- | --- | --- | --- | --- |
|  | IVWM | BC | IVWM | BC |
| TSB | **** | X | **** | X |
| A | X | ns | X | ns |

Liquid forms (the A.D.A.M. method) of REO exhibited significantly higher (p ≤ 0.0001, Dunn’s test. Adjusted p value includes Bonferroni correction) activity in TSB/A and TSB/BC than IVWM/A and IVWM/BC (Figure 8, Table 6, Supplementary Tables S16 and S17). Contrary to the volatile forms, EO’s activity was significantly higher in TSB than in IVWM (p ≤ 0.0001, Dunn’s test. Adjusted p value includes Bonferroni correction). Mean biofilm reductions in these media were 29% and-76%, respectively. No difference was observed for the only-surface dividing condition (A vs BC). TEO completely eradicated *S.aureus* biofilms under all applied growth conditions (Figure 8, Table 6, Supplementary Table S16).

**Table 6.**
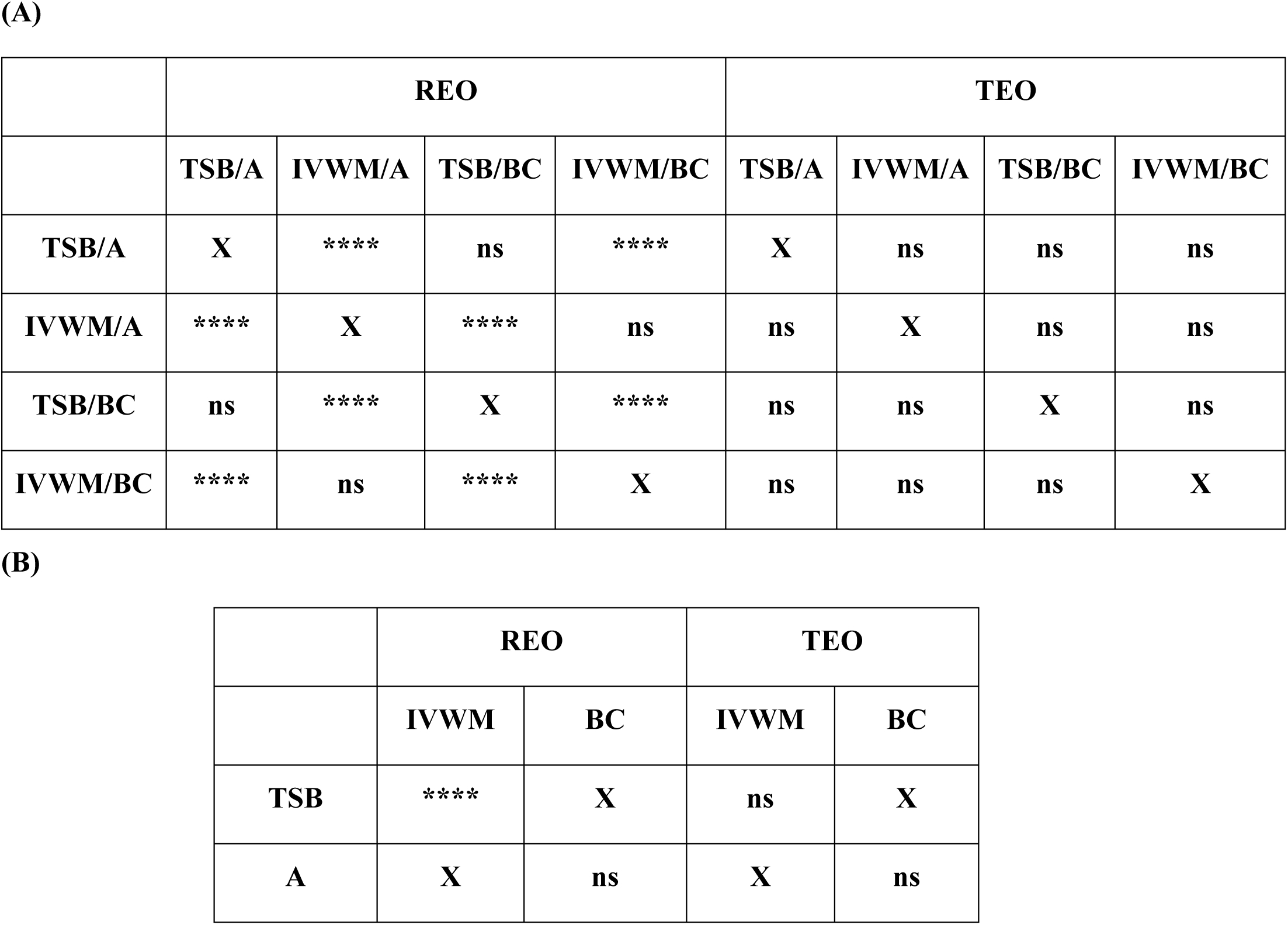
Summarized statistical differences in antibiofilm activity of rosemary essential oil (REO) or thyme essential oil (TEO) against *S. aureus* strains (n=10) cultured on different surfaces: agar (A) or biocellulose (BC) and in different media: tryptic soy broth (TSB) or in in vitro wound milieu (IVWM), assessed with an antibiofilm dressing’s activity measurement method. **(A)** Dividing condition: medium and surface. **(B)** Dividing condition: medium or surface. Dunn’s test was performed. Adjusted p value includes Bonferroni correction. Values of p<0.05 were considered significant, p ≤ 0.0001 was marked with four asterisks. Ns-no significant differences. X-not applicable.

|  | REO |  |  |  | TEO |  |  |  |
| --- | --- | --- | --- | --- | --- | --- | --- | --- |
|  | TSB/A | IVWM/A | TSB/BC | IVWM/BC | TSB/A | IVWM/A | TSB/BC | IVWM/BC |
| TSB/A | X | **** | ns | **** | X | ns | ns | ns |
| IVWM/A | **** | X | **** | ns | ns | X | ns | ns |
| TSB/BC | ns | **** | X | **** | ns | ns | X | ns |
| IVWM/BC | **** | ns | **** | X | ns | ns | ns | X |

(B)
|  | REO |  | TEO |  |
| --- | --- | --- | --- | --- |
|  | IVWM | BC | IVWM | BC |
| TSB | **** | X | ns | X |
| A | X | ns | X | ns |

**Table 7.** Summarized statistical differences in antibiofilm activity of rosemary essential oil (REO) or thyme essential oil (TEO) against *S. aureus* strains (n=10) cultured on different surfaces: agar (A) or biocellulose (BC) and in different media: tryptic soy broth (TSB) or in in vitro wound milieu (IVWM) between the results of an antibiofilm activity of volatile method compounds and an antibiofilm dressing’s activity measurement method. **(A)** Dividing condition: medium and surface. **(B)** Dividing condition: medium or surface. Dunn’s test was performed. Adjusted p value includes Bonferroni correction. Values of p<0.05 were considered significant, p ≤ 0.0001 was marked with four asterisks. Ns-no significant differences. X-not applicable.

|  | REO |  |  |  | TEO |  |  |  |
| --- | --- | --- | --- | --- | --- | --- | --- | --- |
|  | TSB/A | IVWM/A | TSB/BC | IVWM/BC | TSB/A | IVWM/A | TSB/BC | IVWM/BC |
| TSB/A | **** | X | X | X | **** | X | X | X |
| IVWM/A | X | **** | X | X | X | ns | X | X |
| TSB/BC | X | X | **** | X | X | X | **** | X |
| IVWM/BC | X | X | X | **** | X | X | X | ns |

(B)
|  | REO |  |  |  | TEO |  |  |  |
| --- | --- | --- | --- | --- | --- | --- | --- | --- |
|  | TSB | IVWM | A | BC | TSB | IVWM | A | BC |
| TSB | ns | X | X | X | ns | X | X | X |
| IVWM | X | ns | X | X | X | ns | X | X |
| A | X | X | ns | X | X | X | ns | X |
| BC | X | X | X | ns | X | X | X | ns |

Comparison of differences in EOs antibiofilm activity between the ABV and the A.D.A.M. methods revealed statistical significance (p ≤ 0.0001, Dunn’s test. Adjusted p value includes Bonferroni correction) under all four conditions (grouped by medium and surface) in the case of REO. In turn, differences were observed between TSB/A and TSB/BC conditions for TEO (p ≤ 0.0001, Dunn’s test. Adjusted p value includes Bonferroni correction) (Table 7, Supplementary Table S19).

### 3.4 Biofilm visualization

Microscopic visualization was performed to qualitatively assess the impact of REO and TEO volatile fractions on established *S. aureus* biofilms grown on PS in either TSB or IVWM media (Figure 9). Representative strains S3 and S4 were selected for imaging due to their contrasting susceptibility profiles shown via the ABV method.

**Figure 9.**
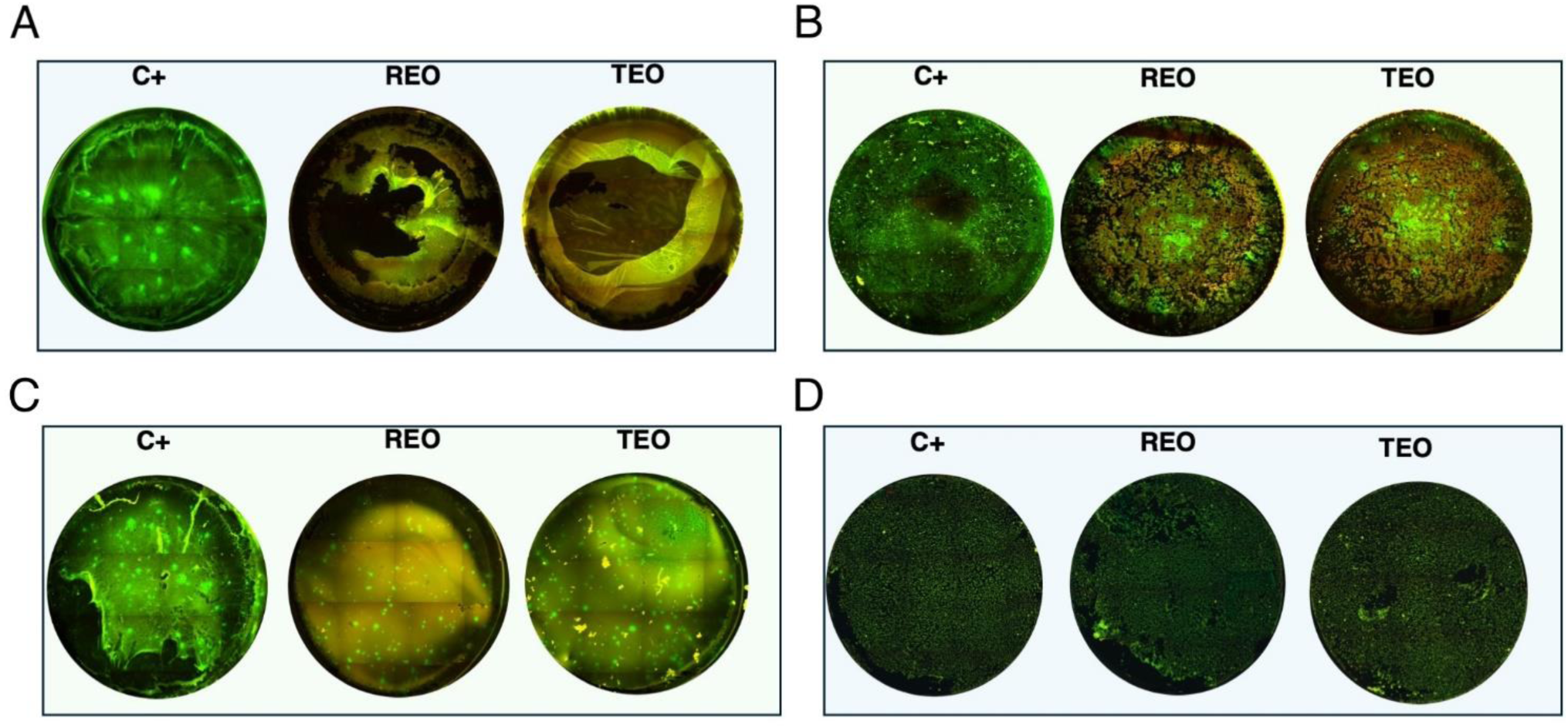
Microscopic visualization of *S. aureus* biofilms (n=2) grown on polystyrene in tryptic soy broth (TSB) or in in vitro wound milieu medium (IVWM) and treated with volatile fractions of rosemary essential oil (REO) or thyme essential oil (TEO). C+ is a growth control setting, where biofilms were non-exposed to antimicrobials. **(A)** S3 strain in TSB. **(B)** S3 strain in IVWM. **(C)** S4 strain in TSB. **(D)** S4 strain in IVWM. All biofilms were stained using the Filmtracer™ LIVE/DEAD™ Biofilm Viability Kit to assess cell viability. Green fluorescence indicates live cells with intact membranes, while red fluorescence marks cells with compromised membrane integrity. Images represent a full-well of a 24-well plate tiling acquired using a 4× objective and ImageJ Software. Microscope Lumascope 620. The well diameter was 15 mm.

The fluorescence staining clearly revealed that both media and EO treatment significantly influenced biofilm architecture and bacterial viability. Even in untreated controls (C+, growth control), differences in biomass density and structure were evident between biofilms grown in TSB and those cultured in IVWM, supporting earlier quantitative findings that nutrient composition affects biofilm development. Upon treatment, both EOs induced visible disruption of the biofilm structure and increased red fluorescence, indicating compromised membrane integrity. However, the extent of this effect varied between oils and was dependent on both the strain and growth conditions. In general, TEO appeared to exert stronger bactericidal and biofilm-disrupting activity than REO, particularly against TSB-grown biofilms, which were thicker and more metabolically active. Conversely, IVWM-grown biofilms exhibited less biomass but appeared more resistant to both EOs, particularly in the case of REO treatment. These findings visually reinforce the notion that essential oil efficacy is highly context-dependent and influenced by strain-level susceptibility, growth medium, and initial biofilm architecture.

### 3.5 Analysis of biofilms’ protein profiles

Analysis with a MALDI-TOF ultrafleXtreme spectrometer was performed to evaluate staphylococcal protein profiles after the biofilms’ exposure to the volatile fractions of REO or TEO. The biofilms were formed on BC and in TSB or IVWM medium (Figure 10). However, the high background of IVWM medium (which is rich in proteins such as fibrinogen, fibronectin and lactoferrin) hindered the detection of bacterial proteins. The following differences between the EOs and C+ in the profile of the S3 strain cultured in TSB were observed: the peak at approximately 4010 m/z was present in the REO and TEO samples but not in the C+. Peaks at about 6230m/z and 3618 m/z were detected only in TEO-treated samples. In the case of the S4 strain in TSB, the mass spectra were obtained only for REO and C+ samples. Five peaks (at 4056 m/z; 4317 m/z; 5045 m/z; 2830 m/z; 8111 m/z) appeared in the spectrum of the bacterial profile after REO treatment. The peak at 6906 m/z was more intense than in the control sample. Four peaks were detected in the C+ spectrum but not in the REO one (at 3014 m/z; 2642 m/z; 2311 m/z; 6228 m/z).

**Figure 10.**
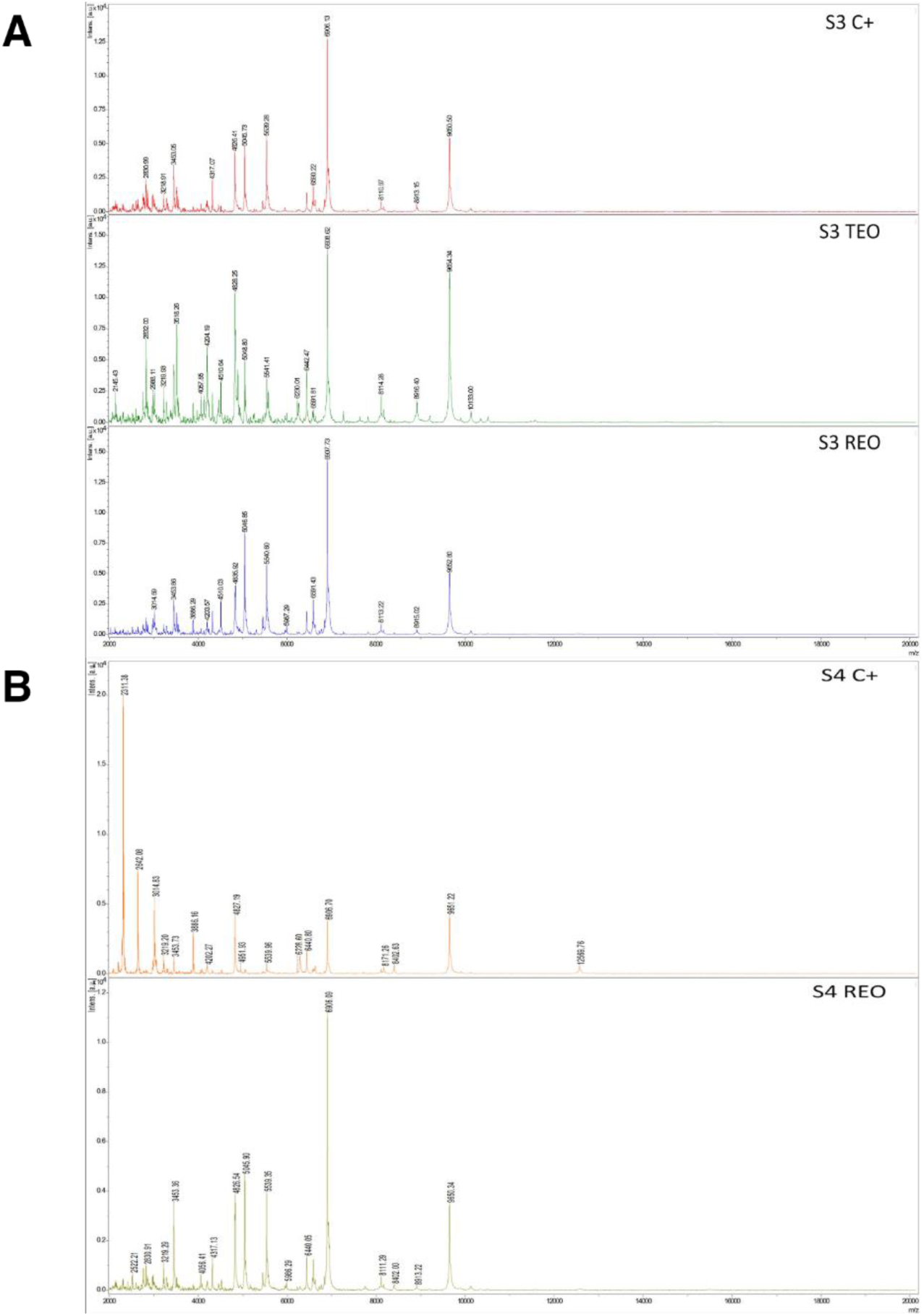
Mass spectra presenting protein profiles of *S. aureus* strains (n=2) cultured on biocellulose in tryptic soy broth after the treatment with volatile fractions of rosemary essential oil (REO), thyme essential oil (TEO), or untreated cells (C+, growth control). **(A)** S3 strain. **(B)** S4 strain. MALDI-TOF (Matrix-Assisted Laser Desorption/Ionization Time-of-Flight Mass Spectrometry) ultrafleXtreme spectrometer.

## 4 Discussion

The concept of Essential Oil Stewardship emerged from our growing awareness of the methodological gaps that limit the translational value of EOs research. Drawing on several years of investigation, our team has developed and progressively refined a framework that guides the selection, evaluation, and interpretation of EOs as anti-infective agents (Brożyna et al., 2020, 2021, 2024). This framework aligns EO testing with the rigor expected of antibiotics or antiseptics and lays the foundation for reproducible and translationally relevant studies.

A key component of this framework is the use of pharmacopoeial-grade EOs, which enhances the relevance of experimental findings. In this study, both rosemary and thyme oils met the quality standards defined by the European Pharmacopoeia XI, with only a minor deviation observed in one constituent. Within the framework, we also employed a panel of *S. aureus* strains, rather than relying on a single reference strain. This approach reflects microbial heterogeneity and improves the applicability of the findings.

Also, biofilm formation was assessed using more than one parameter, i.e. total biomass (cells and extracellular matrix), metabolic activity, and viable cell number. These features were in turn compared across different growth conditions, defined by combinations of two media (TSB and IVWM) and three surface types (polystyrene, biocellulose, and agar polymer). Notably, IVWM simulates the biochemical composition of wound exudate, while biocellulose and agar polymers provide soft, porous surfaces that mimic the topology and structure of wound tissue. To account for different modes of EO delivery, we applied three complementary methods – microtitrate plate method (a.k.a. dilution method), A.D.A.M., and AntiBioVol - allowing us to evaluate EOs’ both contact and vapor-phase activity.

The resulting matrix of variables produced a complex dataset, requiring robust statistical approaches to detect interactions and capture strain-level variability.

The obtained results revealed consistent differences between standard laboratory and infection-relevant settings (Figures 1-3, Tables 1 and 2). Across all evaluated parameters – biomass, metabolic activity and viable cell number – biofilms formed in standard TSB medium were significantly more developed than those cultured in IVWM, regardless of the surface used. TSB promoted robust biofilms with high cell density and metabolic activity. In contrast, biofilms in IVWM were thinner, less metabolically active and composed of fewer viable cells. The IVWM, a formulation mimicking wound exudate, contains host-associated factors such as lactoferrin and lactic acid. These may act as immunological cues, prompting bacteria to organize into smaller, protective clusters, as previously reported (Brożyna et al., 2024). It indicates that the use of IVWM reveals biofilm phenotypes that standard media obscure.

The next analysis on the relationship between biofilm biomass and viable cell number was examined to determine whether total biofilm mass reflects the abundance of living cells. Scatter plot analysis (Figure 5) revealed no strong correlation between these parameters across growth conditions. This indicates that high biomass may result from increased extracellular matrix or dead cell accumulation rather than solely a dense population of viable cells. Similarly, no meaningful correlation was found between biomass and metabolic activity (Figure 4). Not surprisingly, the strongest correlation among the three parameters was observed between metabolic activity and CFU counts (Figure 6), consistent across media and surfaces. Such an observation indicates that metabolic activity is a direct function of both the number of viable cells and their physiological engagement in active processes.

In non-healing wound environments, biofilms may contain large amounts of matrix with only limited viable cells or cells exhibiting dormant metabolic states (Thaarup et al., 2022). Therefore, relying solely on biomass can lead to misinterpretation of biofilm structure and treatment efficacy. The CFU-metabolism correlation highlights these two measures as key endpoints in evaluating EO-mediated antibiofilm effects. The differences in biofilm structure and composition observed under various growth conditions have translational implications. The TSB/PS (polystyrene) setup, characterized by high biomass, high metabolic activity, and large numbers of viable cells, resembles biofilms formed on abiotic medical devices such as catheters, particularly in immunocompromised patients or those with diabetes (Roberts, 2013). In contrast, the IVWM/BC (biocellulose) model produced biofilms with lower metabolic activity and viable cell counts, and a more clustered morphology, consistent with observations from non-healing wound infections (Thaarup et al., 2022).

The inclusion of both simplified (TSB/PS) and clinically-relevant (IVWM/BC) conditions within the same experimental framework enables a more comprehensive understanding of how EOs might perform in clinically distinct environments. It also allows for better prediction of EO efficacy depending on the infection niche - ranging from highly active, surface-associated biofilms to more quiescent, matrix-rich/dormant communities in non-healing wounds. Notably, these two structurally and functionally distinct types of biofilms responded differently to treatment with liquid-phase thyme essential oil (TEO) (Figure 7). Despite the TSB/PS system containing more biomass and viable cells, TEO induced a stronger reduction in both parameters compared to the IVWM/BC setup. This was unexpected, as biofilms with higher biomass, cell density and metabolic activity are generally considered more resistant to antiseptic agents. This result suggests that the presence of robust biofilm in the TSB/PS model does not significantly impair the efficacy of TEO. On the contrary, its activity may be enhanced in metabolically active, cell-dense environments. It also raises the possibility that TEO interferes with bacterial metabolism more efficiently than with structural biofilm components, contributing to greater efficacy in active biofilms. Alternatively, TEO may possess sufficient lipophilicity to penetrate matrix-rich biofilms regardless of density. These findings highlight the need to evaluate EO activity under conditions that reflect both high-burden infections (e.g., contaminated surfaces) and low-metabolism chronic niches (e.g., non-healing wounds), as their performance may differ substantially depending on biofilm architecture and metabolic state. The observed differences in biofilm reduction between TEO and REO (Figure 7) may be partly explained by their distinct chemical compositions and physicochemical interactions with the biofilm matrix. Essential oils are complex mixtures of lipophilic compounds, and their efficacy may be influenced by their ability to partition into hydrophobic biofilm components, such as extracellular polysaccharides and proteins (Stoodley et al., 1999).

Thyme oil, which contains a higher proportion of phenolic compounds like thymol and carvacrol, may interact more strongly with bacterial membranes and disrupt metabolic processes, particularly in biofilms with high cell density and metabolic activity (Kachur and Suntres, 2020). In contrast, rosemary oil, dominated by oxygenated monoterpenes such as 1,8-cineole and α-pinene, may have weaker interactions with dense matrix structures or may require different conditions for optimal activity (Maczka et al., 2021).

The interplay between EO lipophilicity, matrix composition, and bacterial physiology is a promising area for translational research. Understanding these relationships could help match specific EO profiles to infection types, optimizing their use as complementary agents in biofilm-associated infections.

While TEO demonstrated marked differences in antibiofilm efficacy across conditions, REO showed a relatively uniform, though less potent, effect. Its performance appeared largely independent of the growth medium or surface type, suggesting a broader but weaker activity profile. This could indicate a lower sensitivity of REO to environmental variability or a mechanism of action less dependent on specific biofilm phenotypes.

Beyond differences in chemical composition, our results also revealed divergent activity profiles of REO and TEO depending on the testing method. TEO demonstrated higher antibiofilm efficacy in the A.D.A.M. assay, which models gradual diffusion through a viscous matrix, while REO showed relatively better performance in the AntiBioVol assay, designed to assess volatile-phase activity (Figure 8).

These findings indicate that specific EO constituents may differ in potency not only because of differences in the composition/level of antimicrobial substances but also because of differences in their physicochemical behavior - particularly volatility and diffusivity - which in turn affects their bioavailability under different exposure scenarios. For instance, compounds with higher vapor pressure or lower molecular weight may exhibit greater activity in vapor-phase assays, while more hydrophobic or viscous constituents might perform better when directly applied to biofilms in liquid form.

This has important translational implications. Depending on the intended route of administration - topical liquid formulations, impregnated dressings, or vapor-based devices—EOs should be selected and evaluated using methods that reflect their actual clinical deployment. The observed method-dependent differences in REO and TEO activity further underscore the value of employing multiple, complementary models during EO screening.

The integrated design of this study - combining pharmacopoeial-grade oils, clinical strain panels, multiple experimental modalities, and infection-mimicking conditions - demonstrates a pathway toward translating EO research into clinically relevant applications. By aligning experimental models with real-world scenarios (e.g., wound exudate composition, tissue-like surfaces, and polymicrobial complexity), we improve the applicability of *in vitro* findings in real life as well.

Moreover, the method-dependent differences in EO performance, clearly visualized by fluorescence microscopy (Figure 9), highlight the need to match specific EO compositions with suitable clinical delivery forms. Although performed as a proof of concept, the data presented in Figure 10 further demonstrate that exposing the same *S. aureus* strain to two different EOs results in distinct changes in protein expression. Conversely, exposing two different strains to the same EO also leads to different protein expression profiles.

Importantly, the demonstrated strain-level variability in biofilm architecture and EO susceptibility emphasizes the need to move beyond generic claims of EO efficacy. Instead, tailored formulations and targeted indications, guided by systematic preclinical assessment, may allow EOs to complement existing anti-infective strategies, particularly in niches where conventional agents fail or resistance is prevalent.

Essential Oil Stewardship, as proposed here, is not a fixed protocol but a dynamic framework inviting refinement and interdisciplinary input. We hope that this study will encourage broader adoption of rigorous EO testing standards and foster collaboration among microbiologists, pharmacognosists, formulation scientists, and clinicians to responsibly develop EOs as adjuvants in the fight against biofilm-associated infections.

## Conclusions

- Pharmacopoeial-grade essential oils can be systematically evaluated using clinically relevant models.
- Biofilm characteristics are highly dependent on environmental context.
- EO efficacy varies with delivery mode and biofilm environment.
- The Essential Oil Stewardship framework enables reproducible, application-oriented EO testing.

## 6 Author Contributions

Writing-original draft: MB, ZS, KK, BD, AJ. Writing-review and editing: MB, AM, AJ. Conceptualization: MB, AJ. Data curation: MB, ZS. Formal analysis: MB, ZS, AJ. Funding acquisition: MB, AJ. Investigation: MB, KK, BD, AJ. Methodology: MB, ZS, AJ. Project administration: MB, AJ. Software: ZS, MB. Supervision: MB, AJ. Validation: MB, ZS, AJ. Visualization: ZS, MB, AJ.

## 8 Acknowledgments

The Authors would like to express our gratitude to Dr. Weronika Kozłowska from Division of Pharmaceutical Biotechnology, Department of Pharmaceutical Biology and Biotechnology, Wroclaw Medical University, Wroclaw, Poland for the help in the evaluation of essential oils composition.

## 9 Generative AI statement

The authors declare that no Generative AI was used in the creation of this manuscript.

