## Supplementary Material for "Establishing Essential Oil Stewardship Through the Case of Rosemary and Thyme Oils Against *Staphylococcus aureus*"

Supplementary Table S1. Percentage content of particular compounds in **(A)** rosemary essential oil (REO) or **(B)** thyme essential oil (TEO) assessed with GC–MS (gas chromatography-mass spectrometry). EuPh XI ranges- content ranges determined in the European Pharmacopoeia XI. Four main compounds of each EO are bolded. The dashes indicate compounds that were not present in the analyzed EOs. RI - retention index, RT – retention time, SD- standard deviation, ND- not determined in EuPh XI.

**(A)**

| **Compound** | **RI** | **RT** | **Content in REO (%)** | **SD** | **EuPh XI ranges (%)** |
|---|---|---|---|---|---|
| Tricyclene | 921 | 8.81 | 0.41 | 0.01 | ND |
| α-Thujene | 925 | 8.94 | 0.11 | 0.00 | ND |
| α-Pinene | 933 | 9.22 | **21.07** | 0.16 | 18.0-26.0 |
| Camphene | 948 | 9.77 | **8.81** | 0.05 | 8.0-12.0 |
| Thuja-2,4(10)-diene | 952 | 9.90 | 0.30 | 0.01 | ND |
| β-Pinene | 976 | 10.76 | 3.78 | 0.08 | 2.0-6.0 |
| Octen-2-ol/octanone | 984 | 11.04 | 0.20 | 0.01 | ND |
| Myrcene | 989 | 11.20 | 2.70 | 0.05 | 1.5-5.0 |
| α-Phellandrene | 1005 | 11.80 | 0.52 | 0.01 | ND |
| 3-Carene | 1008 | 11.89 | 0.61 | 0.01 | ND |
| α-Terpinene | 1016 | 12.19 | 0.65 | 0.01 | ND |
| p-Cymene | 1024 | 12.48 | 2.46 | 0.04 | 1.0-2.2 |
| Limonene | 1029 | 12.66 | 3.88 | 0.10 | ND |
| 1,8-Cineole | 1032 | 12.79 | **19.98** | 0.11 | 16.0-25.0 |

| | | | | | |
|---|---|---|---|---|---|
| trans-β-Ocimene | 1035 | 12.91 | 0.16 | 0.00 | ND |
| g-Terpinene | 1057 | 13.73 | 1.01 | 0.02 | ND |
| Terpinolene | 1084 | 14.73 | 0.68 | 0.01 | ND |
| Linalool | 1099 | 15.28 | 0.71 | 0.01 | ND |
| Camphor | 1148 | 17.05 | **18.52** | 0.08 | 13.00-21.0 |
| Borneol | 1172 | 17.92 | 3.54 | 0.03 | 2.0-4.5 |
| Terpinen-4-ol | 1180 | 18.23 | 0.68 | 0.01 | ND |
| α-Terpineol | 1194 | 18.76 | 2.32 | 0.02 | 1.0-3.5 |
| Verbenone | 1206 | 19.18 | 1.91 | 0.01 | 0.7-2.5 |
| Bornyl acetate | 1284 | 21.88 | 1.23 | 0.01 | 0.5-2.5 |
| Copaene | 1369 | 24.74 | 0.15 | 0.00 | ND |
| β-Caryophyllene | 1419 | 26.37 | 2.76 | 0.00 | ND |
| α-Humulene | 1456 | 27.50 | 0.50 | 0.00 | ND |
| g-Muurolene | 1479 | 28.23 | 0.11 | 0.00 | ND |
| d-Cadinene | 1517 | 29.41 | 0.12 | 0.02 | ND |

**(B)**

| **Compound** | **RI** | **RT** | **Content in TEO (%)** | **SD** | **EPh XI ranges (%)** |
|---|---|---|---|---|---|
| α-Thujene | 925 | 8.94 | 0.38 | 0.07 | 0.2-1.5 |
| α-Pinene | 932 | 9.19 | 0.91 | 0.16 | ND |
| Camphene | 948 | 9.77 | 1.06 | 0.07 | ND |
| β-Pinene | 976 | 10.75 | 0.31 | 0.02 | ND |
| Myrcene | 988 | 11.19 | 1.11 | 0.09 | 1.0-3.0 |
| α-Terpinene | 1016 | 12.19 | 1.81 | 0.13 | 0.9-2.6 |
| p-Cymene | 1024 | 12.5 | **19.2** | 1.21 | 14.0-28.0 |
| Limonene | 1028 | 12.65 | 0.37 | 0.03 | ND |
| Eucalyptol | 1030 | 12.7 | 0.28 | 0.01 | ND |
| g-Terpinene | 1058 | 13.74 | **9.06** | 0.54 | 4.0-12.0 |
| Linalool | 1099 | 15.28 | 3.21 | 0.55 | 1.5-6.5 |
| Camphor | 1147 | 17.01 | 0.62 | 0.08 | ND |
| Borneol | 1172 | 17.92 | 1.98 | 0.25 | ND |
| Terpinen-4-ol | 1180 | 18.23 | 1.02 | 0.13 | 0.1-2.5 |
| α-Terpineol | 1195 | 18.76 | 0.42 | 0.06 | ND |
| Thymol methyl ether | 1238 | 20.27 | 0.46 | 0.04 | ND |
| Thymol | 1291 | 22.15 | **50.59** | 1.74 | 37.0-55.0 |
| Carvacrol | 1297 | 22.36 | **5.65** | 0.63 | 0.5-5.5 |

| | | | | | |
|---|---|---|---|---|---|
| Caryophyllene | 1426 | 26.37 | 1.66 | 0.11 | ND |
| Carvacrol methyl ether | - | - | - | - | 0.05-1.5 |

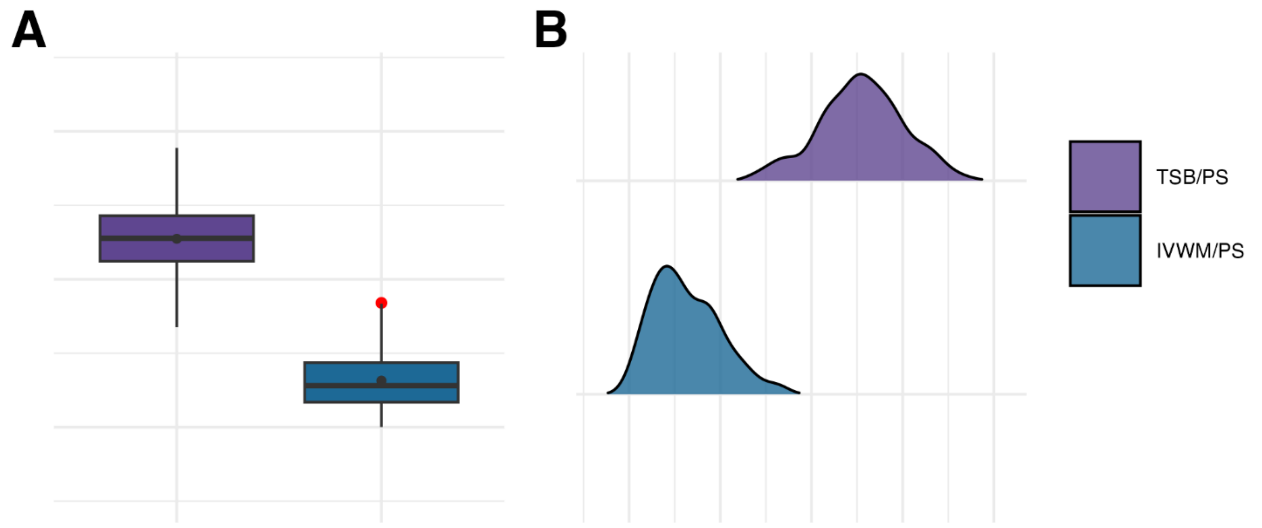


Supplementary Figure S1. **(A, B)** Visual representation of the data distribution for biofilm mass of *S. aureus* strains (n=26) cultured on polystyrene (PS) surface and in different media: tryptic soy broth (TSB) or in in vitro wound milieu (IVWM). Each box displays the interquartile range (IQR; 25th to 75th percentiles), with the bold horizontal line indicating the median. Whiskers extend to the most extreme data points within 1.5 × IQR from the lower and upper quartiles. Mean values are shown as dots.

Supplementary Table S2. Statistical analysis of the data distribution for biofilm mass of *S. aureus* strains (n=26) cultured on polystyrene (PS) surface and in different media: tryptic soy broth (TSB) or in in vitro wound milieu (IVWM). Data was grouped by the following growth conditions: surface and medium. Normal distribution was considered for values of p>0.05. SD- standard deviation, IQR- interquartile range, N- data points.

| Grouping by | Condition | Mean | Median | SD | Min | Max | IQR | N | Skewness | Kurtosis | Shapiro-Wilk p value |
|---|---|---|---|---|---|---|---|---|---|---|---|
| Medium & Surface | TSB/PS | 1.27 | 1.28 | 0.23 | 0.68 | 1.89 | 0.31 | 308 | −0.10 | −0.15 | 0.40 |
| Medium & Surface | IVWM/PS | 0.31 | 0.28 | 0.18 | 0.00 | 0.84 | 0.27 | 298 | 0.60 | −0.19 | 0.00 |

Supplementary Table S3. Parameters of the tested statistic of the differences in biofilm mass of *S. aureus* strains (n=26) cultured on polystyrene (PS) surface and in different media: tryptic soy broth (TSB) or in in vitro wound milieu (IVWM). Data was grouped by the following growth conditions: surface and medium. Dunn's test was performed. Adjusted p value includes Bonferroni correction.

Values of $p<0.05$ were considered significant $p \leq 0.0001$ was marked with four asterisks. N- data points.

| Groupped by | Growth condition 1 | Growth condition 2 | N1 | N2 | mean ranks difference | mean rank1 | mean rank2 | p value | adj. p value | Significance |
|---|---|---|---|---|---|---|---|---|---|---|
| Medium & Surface | IVWM/PS | TSB/PS | 298 | 308 | 302.44 | 149.79 | 452.22 | $2.72 \times 10^{-100}$ | $2.72 \times 10^{-100}$ | **** |

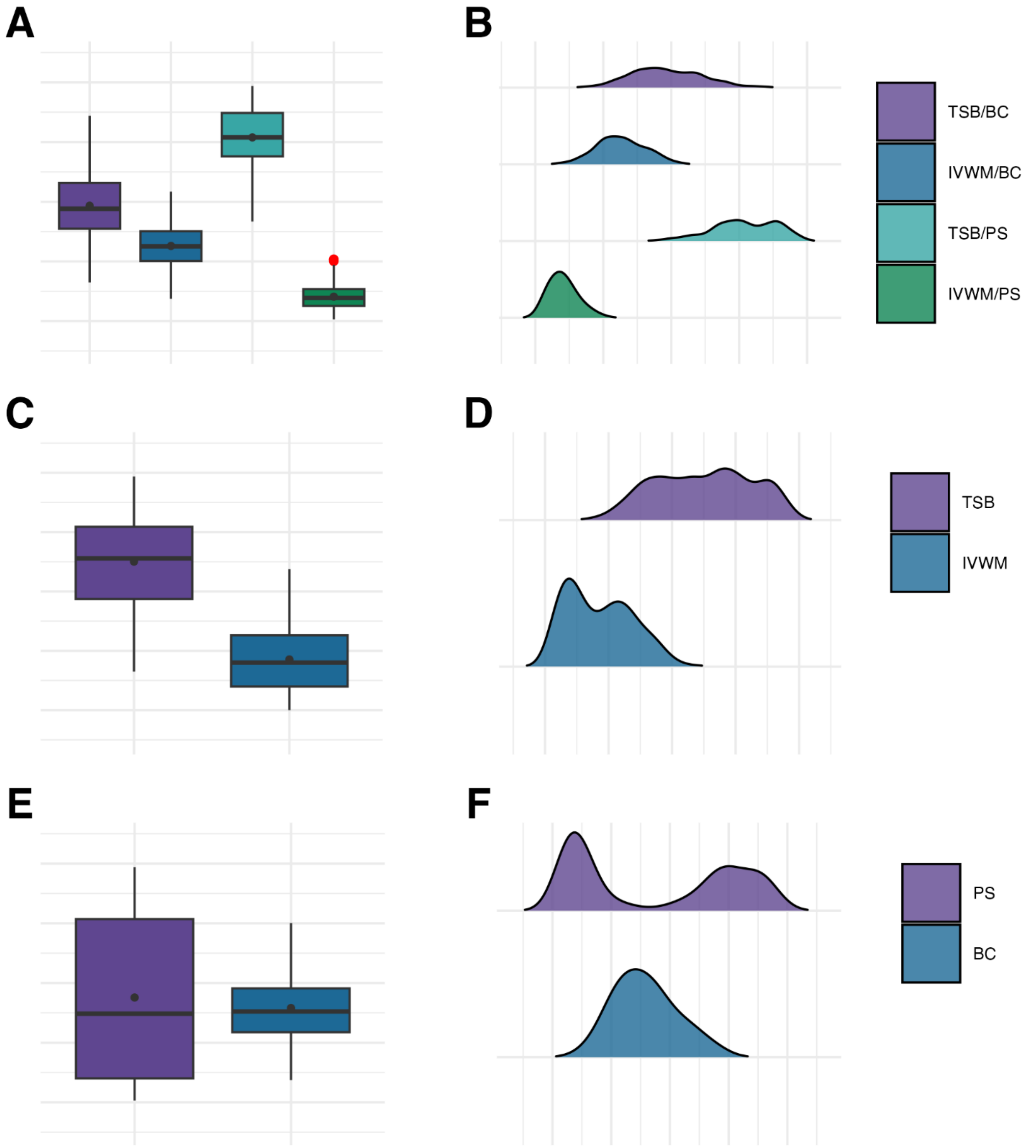

Supplementary Figure S2. Visual representation of the data distribution for biofilm metabolic activity of *S. aureus* strains (n=26) cultured on different surfaces: polystyrene (PS) or biocellulose (BC), and in different media: tryptic soy broth (TSB) or in in vitro wound milieu (IVWM). **(A, B)** Dividing condition: medium and surface. **(C, D)** Dividing condition: medium. **(E, F)** Dividing condition: surface. Each box displays the interquartile range (IQR; 25th to 75th percentiles), with the bold horizontal line indicating the median. Whiskers extend to the most extreme data points within 1.5 × IQR from the lower and upper quartiles. Mean values are shown as dots.

Supplementary Table S4. Statistical analysis of the data distribution for biofilm metabolic activity of *S. aureus* strains (n=26) cultured on different surfaces: polystyrene (PS) or biocellulose (BC), and in different media: tryptic soy broth (TSB) or in in vitro wound milieu (IVWM). Data was grouped by the following growth conditions: surface and medium, or medium only, or surface only. Normal distribution was considered for values of p>0.05. SD- standard deviation, IQR- interquartile range, N- data points.

| Grouping by | Condition | Mean | Median | SD | Min | Max | IQR | N | Skewness | Kurtosis | Shapiro-Wilk p value |
|---|---|---|---|---|---|---|---|---|---|---|---|
| Medium & Surface | TSB/BC | 1.93 | 1.88 | 0.52 | 0.65 | 3.44 | 0.77 | 310 | 0.27 | −0.29 | 0.08 |
| Medium & Surface | IVWM/BC | 1.26 | 1.25 | 0.35 | 0.37 | 2.17 | 0.50 | 309 | 0.09 | −0.40 | 0.28 |
| Medium & Surface | TSB/PS | 3.08 | 3.08 | 0.47 | 1.67 | 3.94 | 0.73 | 309 | −0.37 | −0.43 | 0.00 |
| Medium & Surface | IVWM/PS | 0.41 | 0.39 | 0.21 | 0.03 | 1.03 | 0.28 | 297 | 0.62 | −0.01 | 0.00 |
| Medium | TSB | 2.50 | 2.56 | 0.76 | 0.65 | 3.94 | 1.22 | 624 | −0.08 | −1.00 | 0.00 |
| Medium | IVWM | 0.85 | 0.80 | 0.52 | 0.00 | 2.37 | 0.86 | 623 | 0.39 | −0.85 | 0.00 |
| Surface | PS | 1.76 | 1.49 | 1.37 | 0.03 | 3.94 | 2.67 | 623 | 0.12 | −1.74 | 0.00 |
| Surface | BC | 1.58 | 1.52 | 0.53 | 0.37 | 3.00 | 0.73 | 614 | 0.38 | −0.39 | 0.00 |

Supplementary Table S5. Parameters of the tested statistic of the differences in biofilm metabolic activity of *S. aureus* strains (n=26) cultured on different surfaces: polystyrene (PS) or biocellulose (BC), and in different media: tryptic soy broth (TSB) or in in vitro wound milieu (IVWM). Data was grouped by the following growth conditions: surface and medium, or medium only, or surface only. Kruskal-Wallis test, followed by the Dunn's test, was performed. Adjusted p value includes Bonferroni correction. Only differences classified as statistically significant in ANOVA were included. Values of p<0.05 were considered significant, p ≤ 0.0001 was marked with four asterisks. N- data points.

| Groupped by | Growth condition 1 | Growth condition 2 | N1 | N2 | mean ranks difference | mean rank1 | mean rank2 | p value | adj. p value | Significance |
|---|---|---|---|---|---|---|---|---|---|---|
| Medium & Surface | IVWM/BC | IVWM/PS | 309 | 297 | -335.99 | 491.10 | 155.11 | $1.48 \times 10^{-31}$ | $8.87 \times 10^{-31}$ | **** |
| Medium & Surface | IVWM/BC | TSB/BC | 309 | 310 | 243.98 | 491.10 | 735.07 | $9.56 \times 10^{-18}$ | $5.74 \times 10^{-17}$ | **** |
| Medium & Surface | IVWM/BC | TSB/PS | 309 | 309 | 561.45 | 491.10 | 1052.55 | $1.27 \times 10^{-86}$ | $7.65 \times 10^{-86}$ | **** |
| Medium & Surface | IVWM/PS | TSB/BC | 297 | 310 | 579.96 | 155.11 | 735.07 | $1.19 \times 10^{-90}$ | $7.15 \times 10^{-90}$ | **** |
| Medium & Surface | IVWM/PS | TSB/PS | 297 | 309 | 897.44 | 155.11 | 1052.55 | $6.11 \times 10^{-214}$ | $3.67 \times 10^{-213}$ | **** |
| Medium & Surface | TSB/BC | TSB/PS | 310 | 309 | 317.47 | 735.07 | 1052.55 | $6.16 \times 10^{-29}$ | $3.69 \times 10^{-28}$ | **** |
| Medium | IVWM | TSB | 623 | 624 | 575.48 | 336.03 | 911.51 | $3.82 \times 10^{-175}$ | $3.82 \times 10^{-175}$ | **** |

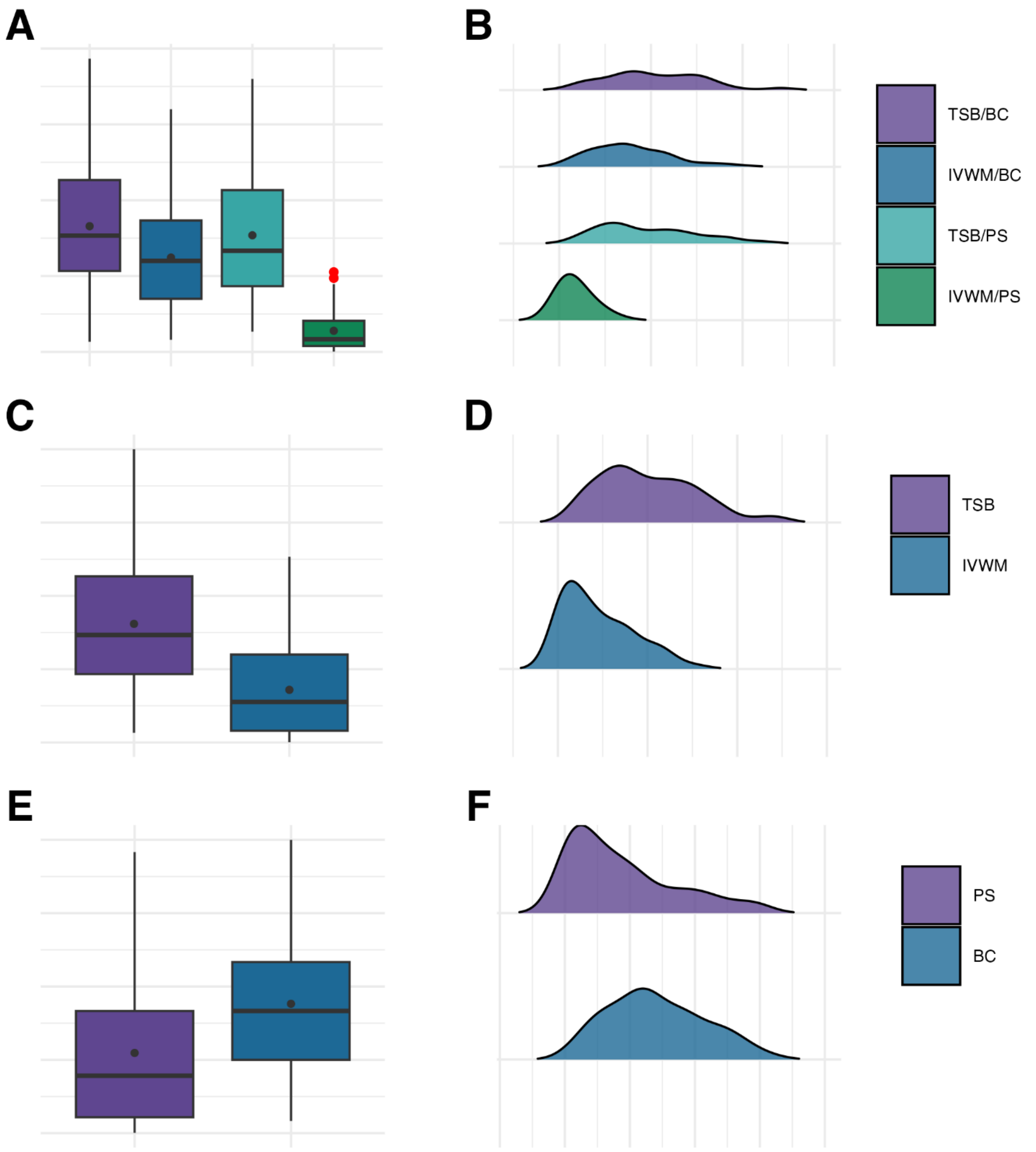


Supplementary Figure S3. Visual representation of the data distribution for biofilm viable cell number of *S. aureus* strains (n=10) cultured on different surfaces: polystyrene (PS) or biocellulose (BC), and in different media: tryptic soy broth (TSB) or in in vitro wound milieu (IVWM). **(A, B)** Dividing condition: medium and surface. **(C, D)** Dividing condition: medium. **(E, F)** Dividing condition: surface. Each box displays the interquartile range (IQR; 25th to 75th percentiles), with the bold horizontal line indicating the median. Whiskers extend to the most extreme data points within 1.5 × IQR from the lower and upper quartiles. Mean values are shown as dots.

Supplementary Table S6. Statistical analysis of the data distribution for biofilm viable cell number of *S. aureus* strains (n=10) cultured on different surfaces: polystyrene (PS) or biocellulose (BC), and in different media: tryptic soy broth (TSB) or in in vitro wound milieu (IVWM). Data was grouped by the following growth conditions: surface and medium, or medium only, or surface only. Normal distribution was considered for values of p>0.05. SD- standard deviation, IQR- interquartile range, N- data points.

| Grouping by | Condition | Mean | Median | SD | Min | Max | IQR | N | Skewness | Kurtosis | Shapiro-Wilk p value |
|---|---|---|---|---|---|---|---|---|---|---|---|
| Medium & Surface | TSB/BC | $4.14 \times 10^{8}$ | $3.83 \times 10^{8}$ | $2.09 \times 10^{8}$ | $3.33 \times 10^{7}$ | $9.67 \times 10^{8}$ | $3.00 \times 10^{8}$ | 60 | 0.41 | −0.18 | 0.15 |
| Medium & Surface | IVWM/BC | $3.11 \times 10^{8}$ | $3.00 \times 10^{8}$ | $1.73 \times 10^{8}$ | $4.00 \times 10^{7}$ | $8.00 \times 10^{8}$ | $2.58 \times 10^{8}$ | 59 | 0.65 | 0.04 | 0.03 |
| Medium & Surface | TSB/PS | $3.84 \times 10^{8}$ | $3.33 \times 10^{8}$ | $2.11 \times 10^{8}$ | $6.67 \times 10^{7}$ | $9.00 \times 10^{8}$ | $3.17 \times 10^{8}$ | 59 | 0.55 | −0.63 | 0.01 |
| Medium & Surface | IVWM/PS | $6.96 \times 10^{7}$ | $4.17 \times 10^{7}$ | $6.61 \times 10^{7}$ | $1.00 \times 10^{6}$ | $2.63 \times 10^{8}$ | $8.32 \times 10^{7}$ | 58 | 1.12 | 0.40 | 0.00 |
| Medium | TSB | $4.05 \times 10^{8}$ | $3.67 \times 10^{8}$ | $2.16 \times 10^{8}$ | $3.33 \times 10^{7}$ | $1.00 \times 10^{9}$ | $3.33 \times 10^{8}$ | 120 | 0.54 | −0.25 | 0.00 |
| Medium | IVWM | $1.80 \times 10^{8}$ | $1.38 \times 10^{8}$ | $1.56 \times 10^{8}$ | $1.00 \times 10^{6}$ | $6.33 \times 10^{8}$ | $2.60 \times 10^{8}$ | 116 | 0.81 | −0.29 | 0.00 |
| Surface | PS | $2.19 \times 10^{8}$ | $1.57 \times 10^{8}$ | $2.05 \times 10^{8}$ | $1.00 \times 10^{6}$ | $7.67 \times 10^{8}$ | $2.90 \times 10^{8}$ | 117 | 0.97 | −0.05 | 0.00 |
| Surface | BC | $3.53 \times 10^{8}$ | $3.33 \times 10^{8}$ | $1.83 \times 10^{8}$ | $3.33 \times 10^{7}$ | $8.00 \times 10^{8}$ | $2.67 \times 10^{8}$ | 117 | 0.32 | −0.72 | 0.02 |

Supplementary Table S7. Parameters of the tested statistic of the differences in biofilm viable cell number of *S. aureus* strains (n=10) cultured on different surfaces: polystyrene (PS) or biocellulose (BC), and in different media: tryptic soy broth (TSB) or in in vitro wound milieu (IVWM). Data was grouped by the following growth conditions: surface and medium, or medium only, or surface only. Kruskal-Wallis test, followed by the Dunn's test, was performed. Adjusted p value includes Bonferroni correction. Only differences classified as statistically significant in ANOVA were included. Values of p<0.05 were considered significant, p ≤ 0.0001 was marked with four asterisks. N- data points, ns- no significant differences assessed in post-hoc tests.

| Groupped by | Growth condition 1 | Growth condition 2 | N1 | N2 | mean ranks difference | mean rank1 | mean rank2 | p value | adj. p value | Significance |
|---|---|---|---|---|---|---|---|---|---|---|
| Medium & Surface | IVWM/BC | IVWM/PS | 59 | 58 | -89.90 | 128.81 | 38.91 | $1.03 \times 10^{-12}$ | $6.20 \times 10^{-12}$ | **** |
| Medium & Surface | IVWM/BC | TSB/BC | 59 | 60 | 28.13 | 128.81 | 156.93 | $2.45 \times 10^{-02}$ | $1.47 \times 10^{-01}$ | ns |
| Medium & Surface | IVWM/BC | TSB/PS | 59 | 59 | 18.55 | 128.81 | 147.36 | $1.40 \times 10^{-01}$ | $8.38 \times 10^{-01}$ | ns |
| Medium & Surface | IVWM/PS | TSB/BC | 58 | 60 | 118.03 | 38.91 | 156.93 | $5.74 \times 10^{-21}$ | $3.45 \times 10^{-20}$ | **** |
| Medium & Surface | IVWM/PS | TSB/PS | 58 | 59 | 108.45 | 38.91 | 147.36 | $8.22 \times 10^{-18}$ | $4.93 \times 10^{-17}$ | **** |
| Medium & Surface | TSB/BC | TSB/PS | 60 | 59 | -9.58 | 156.93 | 147.36 | $4.44 \times 10^{-01}$ | 1.00 | ns |
| Medium | IVWM | TSB | 116 | 120 | 71.78 | 82.00 | 153.78 | $6.46 \times 10^{-16}$ | $6.46 \times 10^{-16}$ | **** |
| Surface | BC | PS | 117 | 117 | -49.74 | 142.37 | 92.63 | $1.86 \times 10^{-08}$ | $1.86 \times 10^{-08}$ | **** |

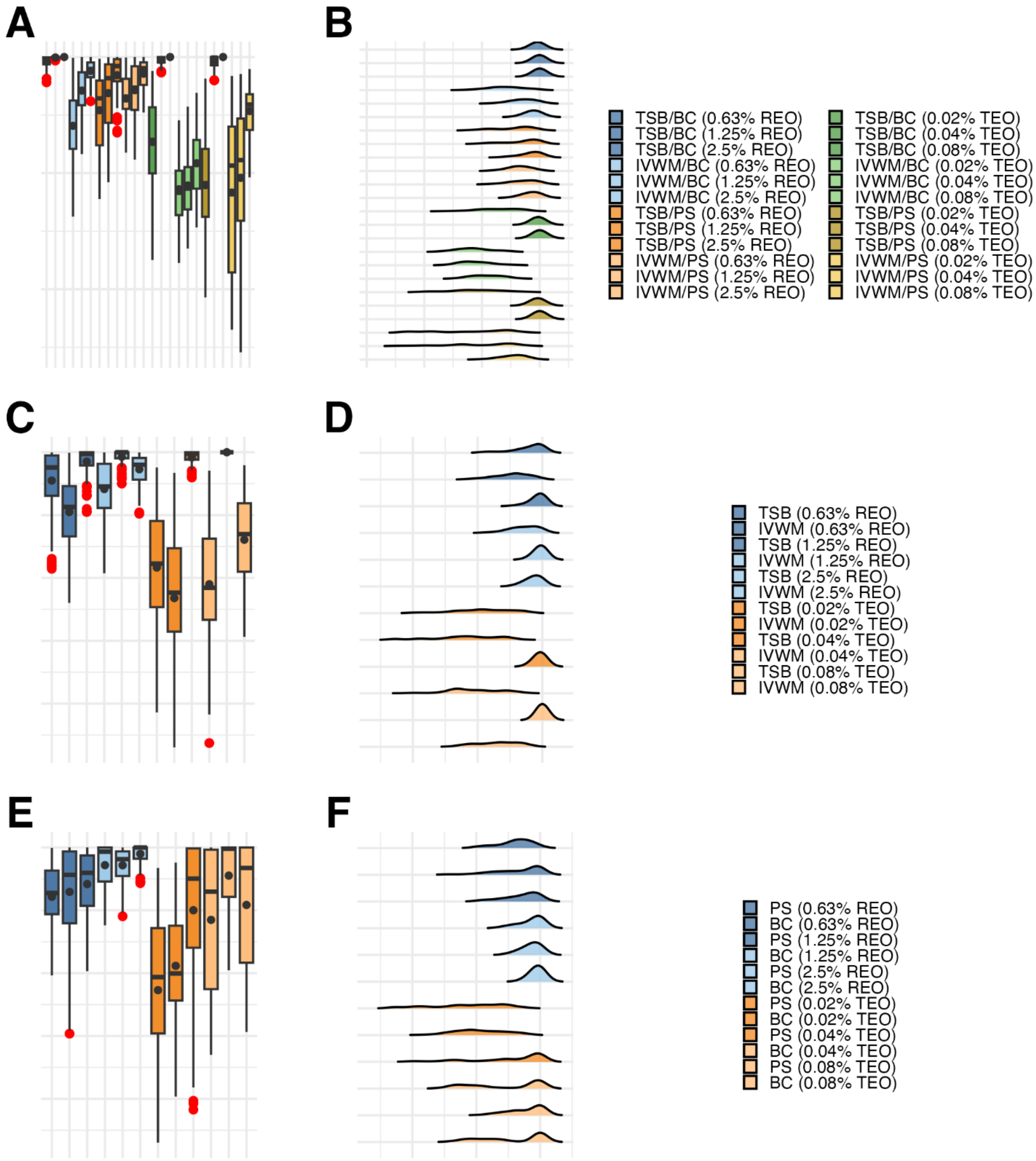


Supplementary Figure S4. Visual representation of the data distribution for antibiofilm activity of rosemary essential oil (REO) or thyme essential oil (TEO) against *S. aureus* strains (n=10) cultured on different surfaces: polystyrene (PS) or biocellulose (BC) and in different media: tryptic soy broth (TSB) or in in vitro wound milieu (IVWM), assessed with a dilution method. **(A, B)** Dividing condition: medium and surface. **(C, D)** Dividing condition: medium. **(E, F)** Dividing condition: surface. Each box displays the interquartile range (IQR; 25th to 75th percentiles), with the bold horizontal line indicating the median. Whiskers extend to the most extreme data points within 1.5 × IQR from the lower and upper quartiles. Mean values are shown as dots.

Supplementary Table S8. Statistical analysis of the data distribution for antibiofilm activity of rosemary essential oil (REO) or thyme essential oil (TEO) against *S. aureus* strains (n=10) cultured on different surfaces: polystyrene (PS) or biocellulose (BC), and in different media: tryptic soy broth (TSB) or in in vitro wound milieu (IVWM), assessed with a dilution method. Data was grouped by the following growth conditions: surface and medium, or medium only, or surface only. Normal distribution was considered for values of $p>0.05$. SD- standard deviation, IQR- interquartile range, N- data points, NaN/ NA- not applicable.

| Grouping by | Condition | Mean | Median | SD | Min | Max | IQR | N | Skewness | Kurtosis | Shapiro-Wilk p value |
|---|---|---|---|---|---|---|---|---|---|---|---|
| Medium & Surface | TSB/BC (0.63% REO) | 95.94 | 97.57 | 5.07 | 78.21 | 100.00 | 6.10 | 51 | −1.60 | 2.23 | 0.00 |
| Medium & Surface | TSB/BC (1.25% REO) | 99.53 | 100.00 | 0.82 | 96.93 | 100.00 | 0.53 | 52 | −1.67 | 1.54 | 0.00 |
| Medium & Surface | TSB/BC (2.5% REO) | 100.00 | 100.00 | 0.00 | 100.00 | 100.00 | 0.00 | 50 | NaN | NaN | NA |
| Medium & Surface | IVWM/BC (0.63% REO) | 40.78 | 41.20 | 30.92 | −37.35 | 99.36 | 41.62 | 57 | −0.13 | −0.51 | 0.75 |
| Medium & Surface | IVWM/BC (1.25% REO) | 71.08 | 72.26 | 20.76 | 16.41 | 100.00 | 28.31 | 56 | −0.50 | −0.42 | 0.03 |
| Medium & Surface | IVWM/BC (2.5% REO) | 88.09 | 89.69 | 9.84 | 61.92 | 100.00 | 12.80 | 54 | −0.76 | −0.26 | 0.00 |
| Medium & Surface | TSB/PS (0.63% REO) | 54.15 | 64.45 | 33.90 | −22.22 | 98.47 | 55.71 | 60 | −0.61 | −0.93 | 0.00 |
| Medium & Surface | TSB/PS (1.25% REO) | 68.90 | 81.78 | 31.52 | −7.76 | 99.79 | 50.36 | 60 | −0.92 | −0.52 | 0.00 |
| Medium & Surface | TSB/PS (2.5% REO) | 84.29 | 92.23 | 17.59 | 34.89 | 100.00 | 19.10 | 57 | −1.19 | 0.43 | 0.00 |
| Medium & Surface | IVWM/PS (0.63% REO) | 64.48 | 62.84 | 18.45 | 17.72 | 99.62 | 25.85 | 58 | −0.35 | −0.29 | 0.19 |
| Medium & Surface | IVWM/PS (1.25% REO) | 71.51 | 73.66 | 23.22 | 11.73 | 100.00 | 37.63 | 60 | −0.62 | −0.47 | 0.00 |
| Medium & Surface | IVWM/PS (2.5% REO) | 85.63 | 90.29 | 12.65 | 52.18 | 100.00 | 19.45 | 60 | −0.67 | −0.57 | 0.00 |

| Grouping by | Condition | Mean | Median | SD | Min | Max | IQR | N | Skewness | Kurtosis | Shapiro-Wilk p value |
|---|---|---|---|---|---|---|---|---|---|---|---|
| Medium & Surface | TSB/BC (0.02% TEO) | 26.91 | 28.42 | 37.03 | −74.74 | 87.93 | 56.96 | 60 | −0.48 | −0.40 | 0.11 |
| Medium & Surface | TSB/BC (0.04% TEO) | 97.27 | 98.80 | 3.60 | 86.63 | 100.00 | 4.79 | 53 | −1.35 | 0.90 | 0.00 |
| Medium & Surface | TSB/BC (0.08% TEO) | 100.00 | 100.00 | 0.00 | 100.00 | 100.00 | 0.00 | 56 | NaN | NaN | NA |
| Medium & Surface | IVWM/BC (0.02% TEO) | −13.86 | −16.89 | 29.15 | −77.25 | 45.51 | 38.24 | 59 | 0.27 | −0.57 | 0.28 |
| Medium & Surface | IVWM/BC (0.04% TEO) | −10.64 | −13.24 | 27.08 | −64.99 | 57.25 | 36.54 | 60 | 0.53 | −0.24 | 0.07 |
| Medium & Surface | IVWM/BC (0.08% TEO) | 8.52 | 8.20 | 26.20 | −46.81 | 69.14 | 41.98 | 60 | 0.06 | −0.88 | 0.43 |
| Medium & Surface | TSB/PS (0.02% TEO) | −10.32 | −8.75 | 46.53 | −106.91 | 81.44 | 58.46 | 60 | −0.14 | −0.65 | 0.55 |
| Medium & Surface | TSB/PS (0.04% TEO) | 96.02 | 99.46 | 5.63 | 79.96 | 100.00 | 7.08 | 56 | −1.46 | 1.32 | 0.00 |
| Medium & Surface | TSB/PS (0.08% TEO) | 100.00 | 100.00 | 0.00 | 100.00 | 100.00 | 0.00 | 51 | NaN | NaN | NA |
| Medium & Surface | IVWM/PS (0.02% TEO) | −16.72 | 6.53 | 66.83 | −134.82 | 83.66 | 125.54 | 60 | −0.44 | −1.29 | 0.00 |
| Medium & Surface | IVWM/PS (0.04% TEO) | −4.00 | 11.26 | 60.40 | −154.24 | 85.30 | 87.94 | 60 | −0.62 | −0.61 | 0.00 |
| Medium & Surface | IVWM/PS (0.08% TEO) | 53.51 | 59.04 | 21.34 | −3.65 | 89.61 | 30.78 | 59 | −0.66 | −0.16 | 0.02 |

| Grouping by | Condition | Mean | Median | SD | Min | Max | IQR | N | Skewness | Kurtosis | Shapiro-Wilk p value |
|---|---|---|---|---|---|---|---|---|---|---|---|
| Medium | TSB (0.63% REO) | 77.48 | 87.91 | 25.41 | 7.29 | 100.00 | 32.12 | 113 | −1.27 | 0.68 | 0.00 |
| Medium | IVWM (0.63% REO) | 52.47 | 56.47 | 27.74 | −19.88 | 99.62 | 40.08 | 116 | −0.51 | −0.36 | 0.00 |
| Medium | TSB (1.25% REO) | 92.61 | 98.69 | 11.35 | 52.45 | 100.00 | 10.53 | 104 | −1.83 | 2.77 | 0.00 |
| Medium | IVWM (1.25% REO) | 70.72 | 72.77 | 22.75 | 3.75 | 100.00 | 35.10 | 117 | −0.67 | −0.15 | 0.00 |
| Medium | TSB (2.5% REO) | 96.41 | 100.00 | 6.27 | 75.08 | 100.00 | 4.78 | 104 | −1.81 | 2.22 | 0.00 |
| Medium | IVWM (2.5% REO) | 86.48 | 90.08 | 11.85 | 51.06 | 100.00 | 17.26 | 115 | −0.86 | 0.02 | 0.00 |
| Medium | TSB (0.02% TEO) | 8.30 | 11.38 | 45.86 | −106.91 | 87.93 | 68.58 | 120 | −0.43 | −0.44 | 0.01 |
| Medium | IVWM (0.02% TEO) | −15.99 | −11.68 | 51.84 | −134.82 | 83.66 | 66.37 | 120 | −0.50 | −0.44 | 0.00 |
| Medium | TSB (0.04% TEO) | 95.89 | 98.80 | 5.62 | 79.96 | 100.00 | 5.87 | 115 | −1.44 | 1.10 | 0.00 |
| Medium | IVWM (0.04% TEO) | −4.99 | −7.76 | 43.48 | −131.36 | 85.30 | 64.56 | 118 | −0.22 | −0.22 | 0.04 |
| Medium | TSB (0.08% TEO) | 100.00 | 100.00 | 0.00 | 100.00 | 100.00 | 0.00 | 107 | NaN | NaN | NA |
| Medium | IVWM (0.08% TEO) | 30.42 | 34.89 | 32.99 | −46.81 | 89.61 | 54.49 | 120 | −0.27 | −0.95 | 0.00 |

| Grouping by | Condition | Mean | Median | SD | Min | Max | IQR | N | Skewness | Kurtosis | Shapiro-Wilk p value |
|---|---|---|---|---|---|---|---|---|---|---|---|
| Surface | PS (0.63% REO) | 60.72 | 63.74 | 25.67 | −1.83 | 99.62 | 34.68 | 116 | −0.74 | −0.27 | 0.00 |
| Surface | BC (0.63% REO) | 64.65 | 78.21 | 37.20 | −48.13 | 100.00 | 57.15 | 119 | −0.97 | 0.15 | 0.00 |
| Surface | PS (1.25% REO) | 70.86 | 79.80 | 26.76 | 1.55 | 100.00 | 40.04 | 119 | −0.87 | −0.27 | 0.00 |
| Surface | BC (1.25% REO) | 85.90 | 96.64 | 18.39 | 37.99 | 100.00 | 27.09 | 114 | −1.11 | −0.10 | 0.00 |
| Surface | PS (2.5% REO) | 85.83 | 90.68 | 13.86 | 45.26 | 100.00 | 19.66 | 115 | −0.93 | −0.07 | 0.00 |
| Surface | BC (2.5% REO) | 94.92 | 100.00 | 7.49 | 71.60 | 100.00 | 9.02 | 110 | −1.43 | 1.06 | 0.00 |
| Surface | PS (0.02% TEO) | −13.52 | −3.14 | 57.43 | −134.82 | 83.66 | 83.81 | 120 | −0.44 | −0.84 | 0.00 |
| Surface | BC (0.02% TEO) | 5.82 | −0.23 | 40.02 | −98.15 | 87.93 | 59.53 | 120 | 0.02 | −0.69 | 0.24 |
| Surface | PS (0.04% TEO) | 50.05 | 75.21 | 59.01 | −108.61 | 100.00 | 78.99 | 117 | −1.08 | −0.02 | 0.00 |
| Surface | BC (0.04% TEO) | 42.37 | 64.83 | 56.72 | −64.99 | 100.00 | 110.91 | 120 | −0.27 | −1.66 | 0.00 |
| Surface | PS (0.08% TEO) | 77.60 | 98.91 | 26.79 | 2.29 | 100.00 | 39.31 | 118 | −0.84 | −0.46 | 0.00 |
| Surface | BC (0.08% TEO) | 54.24 | 83.69 | 49.48 | −46.81 | 100.00 | 91.72 | 120 | −0.38 | −1.50 | 0.00 |

Supplementary Table S9. Parameters of the tested statistic of the differences for antibiofilm activity of rosemary essential oil against *S. aureus* strains (n=10) cultured on different surfaces: polystyrene (PS) or biocellulose (BC), and in different media: tryptic soy broth (TSB) or in in vitro wound milieu (IVWM), assessed with a dilution method. Data was grouped by the following growth conditions: surface and medium, or medium only, or surface only. Kruskal-Wallis test, followed by the Dunn's test, was performed. Adjusted p value includes Bonferroni correction. Only differences classified as statistically significant in ANOVA were included. Values of $p<0.05$ were considered significant, $p \leq 0.05$ was marked with one asterisk, $p \leq 0.01$ was marked with two asterisks, $p \leq 0.0001$ was marked with four asterisks. N- data points, ns- no significant differences assessed in post-hoc tests.

| Groupped by | concentration | Growth condition 1 | Growth condition 2 | N1 | N2 | mean ranks difference | mean rank1 | mean rank2 | p value | adj. p value | Significance |
|---|---|---|---|---|---|---|---|---|---|---|---|
| Medium & Surface | 2.50 | IVWM/BC | IVWM/PS | 53 | 60 | -5.90 | 87.70 | 81.80 | $6.10 \times 10^{-01}$ | 1.00 | ns |
| Medium & Surface | 2.50 | IVWM/BC | TSB/BC | 53 | 50 | 92.80 | 87.70 | 180.50 | $1.65 \times 10^{-14}$ | $9.91 \times 10^{-14}$ | **** |
| Medium & Surface | 2.50 | IVWM/BC | TSB/PS | 53 | 53 | 3.91 | 87.70 | 91.60 | $7.43 \times 10^{-01}$ | 1.00 | ns |
| Medium & Surface | 2.50 | IVWM/PS | TSB/BC | 60 | 50 | 98.70 | 81.80 | 180.50 | $4.30 \times 10^{-17}$ | $2.58 \times 10^{-16}$ | **** |
| Medium & Surface | 2.50 | IVWM/PS | TSB/PS | 60 | 53 | 9.80 | 81.80 | 91.60 | $3.96 \times 10^{-01}$ | 1.00 | ns |
| Medium & Surface | 2.50 | TSB/BC | TSB/PS | 50 | 53 | -88.90 | 180.50 | 91.60 | $1.95 \times 10^{-13}$ | $1.17 \times 10^{-12}$ | **** |
| Medium & Surface | 1.25 | IVWM/BC | IVWM/PS | 56 | 60 | 4.37 | 86.77 | 91.13 | $7.11 \times 10^{-01}$ | 1.00 | ns |
| Medium & Surface | 1.25 | IVWM/BC | TSB/BC | 56 | 44 | 106.32 | 86.77 | 193.09 | $9.34 \times 10^{-17}$ | $5.60 \times 10^{-16}$ | **** |
| Medium & Surface | 1.25 | IVWM/BC | TSB/PS | 56 | 60 | 4.68 | 86.77 | 91.45 | $6.91 \times 10^{-01}$ | 1.00 | ns |
| Medium & Surface | 1.25 | IVWM/PS | TSB/BC | 60 | 44 | 101.96 | 91.13 | 193.09 | $5.91 \times 10^{-16}$ | $3.55 \times 10^{-15}$ | **** |
| Medium & Surface | 1.25 | IVWM/PS | TSB/PS | 60 | 60 | 0.32 | 91.13 | 91.45 | $9.78 \times 10^{-01}$ | 1.00 | ns |
| Medium & Surface | 1.25 | TSB/BC | TSB/PS | 44 | 60 | -101.64 | 193.09 | 91.45 | $7.26 \times 10^{-16}$ | $4.36 \times 10^{-15}$ | **** |
| Medium & Surface | 0.63 | IVWM/BC | IVWM/PS | 57 | 58 | 37.64 | 67.67 | 105.31 | $1.84 \times 10^{-03}$ | $1.10 \times 10^{-02}$ | * |
| Medium & Surface | 0.63 | IVWM/BC | TSB/BC | 57 | 49 | 127.23 | 67.67 | 194.90 | $6.88 \times 10^{-24}$ | $4.13 \times 10^{-23}$ | **** |
| Medium & Surface | 0.63 | IVWM/BC | TSB/PS | 57 | 60 | 27.08 | 67.67 | 94.75 | $2.38 \times 10^{-02}$ | $1.43 \times 10^{-01}$ | ns |
| Medium & Surface | 0.63 | IVWM/PS | TSB/BC | 58 | 49 | 89.59 | 105.31 | 194.90 | $1.04 \times 10^{-12}$ | $6.24 \times 10^{-12}$ | **** |
| Medium & Surface | 0.63 | IVWM/PS | TSB/PS | 58 | 60 | -10.56 | 105.31 | 94.75 | $3.76 \times 10^{-01}$ | 1.00 | ns |
| Medium & Surface | 0.63 | TSB/BC | TSB/PS | 49 | 60 | -100.15 | 194.90 | 94.75 | $1.00 \times 10^{-15}$ | $6.02 \times 10^{-15}$ | **** |
| Medium | 2.50 | IVWM | TSB | 113 | 103 | 50.19 | 84.57 | 134.76 | $1.89 \times 10^{-09}$ | $1.89 \times 10^{-09}$ | **** |
| Medium | 1.25 | IVWM | TSB | 116 | 104 | 45.43 | 89.03 | 134.45 | $1.17 \times 10^{-07}$ | $1.17 \times 10^{-07}$ | **** |
| Medium | 0.63 | IVWM | TSB | 115 | 109 | 53.12 | 86.65 | 139.77 | $8.68 \times 10^{-10}$ | $8.68 \times 10^{-10}$ | **** |
| Surface | 2.50 | BC | PS | 103 | 113 | -46.35 | 132.75 | 86.40 | $2.90 \times 10^{-08}$ | $2.90 \times 10^{-08}$ | **** |
| Surface | 1.25 | BC | PS | 100 | 120 | -42.26 | 133.55 | 91.29 | $8.84 \times 10^{-07}$ | $8.84 \times 10^{-07}$ | **** |
| Surface | 0.63 | BC | PS | 106 | 118 | -26.54 | 126.48 | 99.94 | $2.21 \times 10^{-03}$ | $2.21 \times 10^{-03}$ | ** |

Supplementary Table S10. Parameters of the tested statistic of the differences for antibiofilm activity of thyme essential oil against *S. aureus* strains (n=10) cultured on different surfaces: polystyrene (PS) or biocellulose (BC), and in different media: tryptic soy broth (TSB) or in in vitro wound milieu (IVWM), assessed with a dilution method. Data was grouped by the following growth conditions: surface and medium, or medium only, or surface only. Kruskal-Wallis test, followed by the Dunn's test, was performed. Adjusted p value includes Bonferroni correction. Only differences classified as statistically significant in ANOVA were included. Values of $p<0.05$ were considered significant, $p \leq 0.05$ was marked with one asterisk, $p \leq 0.01$ was marked with two asterisks, $p \leq 0.001$ was marked with three asterisks, $p \leq 0.0001$ was marked with four asterisks. N- data points, ns- no significant differences assessed in post-hoc tests.

| Groupped by | concentration | Growth condition 1 | Growth condition 2 | N1 | N2 | mean ranks difference | mean rank1 | mean rank2 | p value | adj. p value | Significance |
|---|---|---|---|---|---|---|---|---|---|---|---|
| Medium & Surface | 0.08 | IVWM/BC | IVWM/PS | 60 | 59 | 47.40 | 36.50 | 83.90 | $2.89 \times 10^{-05}$ | $1.73 \times 10^{-04}$ | *** |
| Medium & Surface | 0.08 | IVWM/BC | TSB/BC | 60 | 56 | 136.50 | 36.50 | 173.00 | $1.44 \times 10^{-32}$ | $8.62 \times 10^{-32}$ | **** |
| Medium & Surface | 0.08 | IVWM/BC | TSB/PS | 60 | 51 | 136.50 | 36.50 | 173.00 | $4.44 \times 10^{-31}$ | $2.67 \times 10^{-30}$ | **** |
| Medium & Surface | 0.08 | IVWM/PS | TSB/BC | 59 | 56 | 89.10 | 83.90 | 173.00 | $1.11 \times 10^{-14}$ | $6.67 \times 10^{-14}$ | **** |
| Medium & Surface | 0.08 | IVWM/PS | TSB/PS | 59 | 51 | 89.10 | 83.90 | 173.00 | $4.75 \times 10^{-14}$ | $2.85 \times 10^{-13}$ | **** |
| Medium & Surface | 0.08 | TSB/BC | TSB/PS | 56 | 51 | 0.00 | 173.00 | 173.00 | 1.00 | 1.00 | ns |
| Medium & Surface | 0.04 | IVWM/BC | IVWM/PS | 60 | 60 | 8.62 | 56.20 | 64.82 | $4.65 \times 10^{-01}$ | 1.00 | ns |
| Medium & Surface | 0.04 | IVWM/BC | TSB/BC | 60 | 51 | 116.78 | 56.20 | 172.98 | $2.30 \times 10^{-21}$ | $1.38 \times 10^{-20}$ | **** |
| Medium & Surface | 0.04 | IVWM/BC | TSB/PS | 60 | 53 | 115.82 | 56.20 | 172.02 | $1.91 \times 10^{-21}$ | $1.15 \times 10^{-20}$ | **** |
| Medium & Surface | 0.04 | IVWM/PS | TSB/BC | 60 | 51 | 108.16 | 64.82 | 172.98 | $1.49 \times 10^{-18}$ | $8.97 \times 10^{-18}$ | **** |
| Medium & Surface | 0.04 | IVWM/PS | TSB/PS | 60 | 53 | 107.20 | 64.82 | 172.02 | $1.34 \times 10^{-18}$ | $8.05 \times 10^{-18}$ | **** |
| Medium & Surface | 0.04 | TSB/BC | TSB/PS | 51 | 53 | -0.96 | 172.98 | 172.02 | $9.40 \times 10^{-01}$ | 1.00 | ns |
| Medium & Surface | 0.02 | IVWM/BC | IVWM/PS | 59 | 60 | 15.96 | 96.34 | 112.30 | $2.08 \times 10^{-01}$ | 1.00 | ns |
| Medium & Surface | 0.02 | IVWM/BC | TSB/BC | 59 | 60 | 66.83 | 96.34 | 163.17 | $1.35 \times 10^{-07}$ | $8.10 \times 10^{-07}$ | **** |
| Medium & Surface | 0.02 | IVWM/BC | TSB/PS | 59 | 60 | 11.46 | 96.34 | 107.80 | $3.66 \times 10^{-01}$ | 1.00 | ns |
| Medium & Surface | 0.02 | IVWM/PS | TSB/BC | 60 | 60 | 50.87 | 112.30 | 163.17 | $5.58 \times 10^{-05}$ | $3.35 \times 10^{-04}$ | *** |
| Medium & Surface | 0.02 | IVWM/PS | TSB/PS | 60 | 60 | -4.50 | 112.30 | 107.80 | $7.21 \times 10^{-01}$ | 1.00 | ns |
| Medium & Surface | 0.02 | TSB/BC | TSB/PS | 60 | 60 | -55.37 | 163.17 | 107.80 | $1.15 \times 10^{-05}$ | $6.92 \times 10^{-05}$ | **** |
| Medium | 0.08 | IVWM | TSB | 119 | 107 | 113.00 | 60.00 | 173.00 | $7.64 \times 10^{-43}$ | $7.64 \times 10^{-43}$ | **** |
| Medium | 0.04 | IVWM | TSB | 120 | 104 | 111.98 | 60.51 | 172.49 | $2.77 \times 10^{-38}$ | $2.77 \times 10^{-38}$ | **** |
| Medium | 0.02 | IVWM | TSB | 119 | 120 | 31.10 | 104.39 | 135.48 | $5.08 \times 10^{-04}$ | $5.08 \times 10^{-04}$ | *** |
| Surface | 0.08 | BC | PS | 116 | 110 | 22.81 | 102.40 | 125.21 | $5.56 \times 10^{-03}$ | $5.56 \times 10^{-03}$ | ** |
| Surface | 0.02 | BC | PS | 119 | 120 | -19.98 | 130.03 | 110.05 | $2.55 \times 10^{-02}$ | $2.55 \times 10^{-02}$ | * |

Supplementary Table S11. Parameters of the tested statistic of the differences between antibiofilm activity of rosemary essential oil (REO) and thyme essential oil (TEO) against *S. aureus* strains (n=10) cultured on different surfaces: polystyrene (PS) or biocellulose (BC), and in different media: tryptic soy broth (TSB) or in in vitro wound milieu (IVWM), assessed with a dilution method. Data was grouped by the following growth conditions: surface and medium, or medium only, or surface only. Kruskal-Wallis test, followed by the Dunn's test, was performed. Adjusted p value includes Bonferroni correction. Only differences classified as statistically significant in ANOVA were included. Values of $p<0.05$ were considered significant, $p \leq 0.001$ was marked with three asterisks, $p \leq 0.0001$ was marked with four asterisks. N- data points.

| Groupped by | Growth condition | Essential oil1 | Essential oil2 | N1 | N2 | mean ranks difference | mean rank1 | mean rank2 | p value | adj. p value | Significance |
|---|---|---|---|---|---|---|---|---|---|---|---|
| Medium & Surface | TSB/BC | REO | TEO | 143 | 167 | -52.94 | 184.02 | 131.08 | $1.41 \times 10^{-08}$ | $1.41 \times 10^{-08}$ | **** |
| Medium & Surface | IVWM/BC | REO | TEO | 166 | 179 | -153.20 | 252.49 | 99.28 | $4.14 \times 10^{-46}$ | $4.14 \times 10^{-46}$ | **** |
| Medium & Surface | TSB/PS | REO | TEO | 173 | 164 | 37.93 | 150.54 | 188.47 | $3.22 \times 10^{-04}$ | $3.22 \times 10^{-04}$ | *** |
| Medium & Surface | IVWM/PS | REO | TEO | 178 | 179 | -134.26 | 246.32 | 112.06 | $1.01 \times 10^{-34}$ | $1.01 \times 10^{-34}$ | **** |

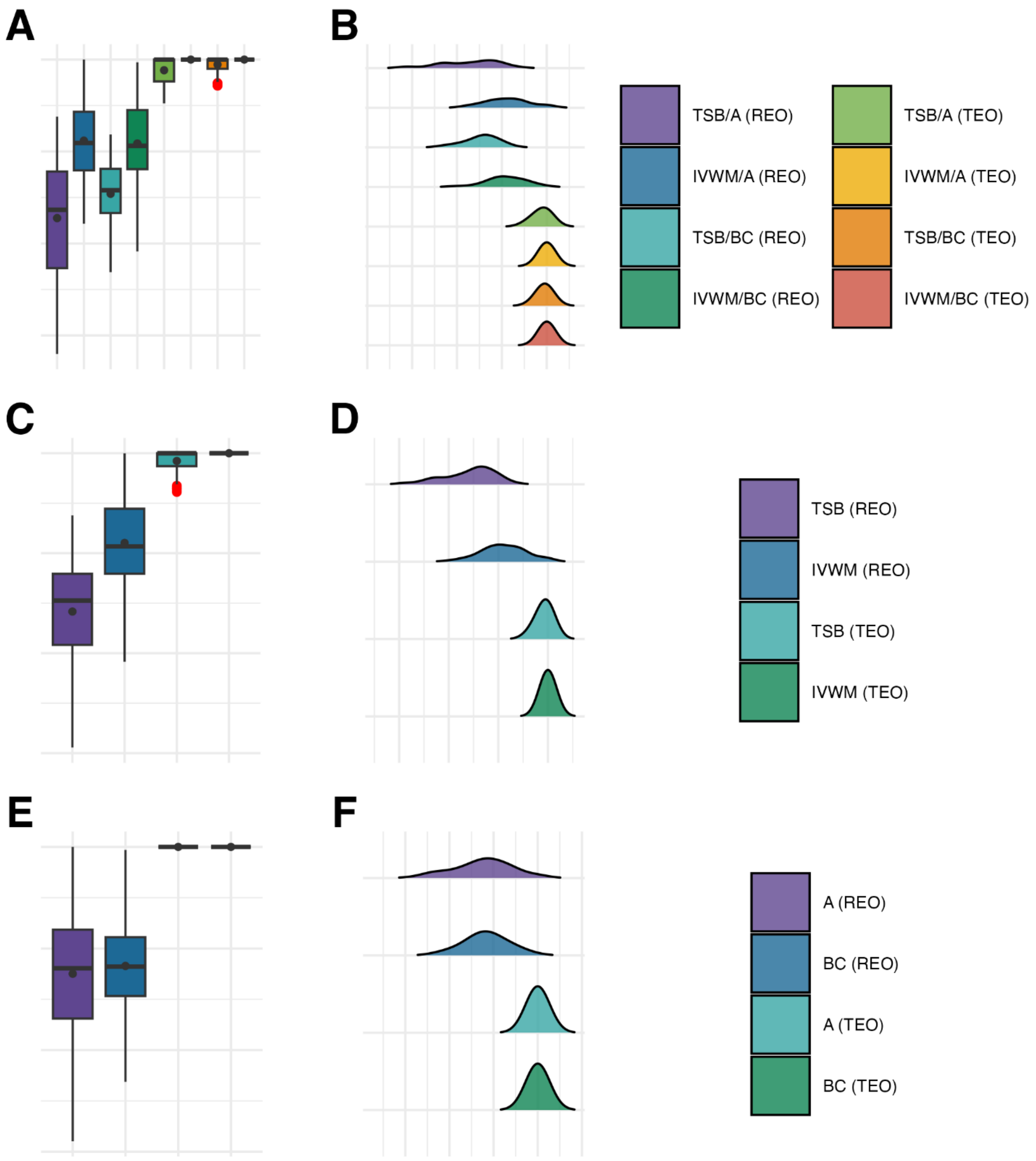


Supplementary Figure S5. Visual representation of the data distribution for antibiofilm activity of rosemary essential oil (REO) or thyme essential oil (TEO) against *S. aureus* strains (n=10) cultured on different surfaces: agar (A) or biocellulose (BC) and in different media: tryptic soy broth (TSB) or in in vitro wound milieu (IVWM), assessed with an antibiofilm activity of volatile compounds method. **(A, B)** Dividing condition: medium and surface. **(C, D)** Dividing condition: medium. **(E, F)** Dividing condition: surface. Each box displays the interquartile range (IQR; 25th to 75th percentiles), with the bold horizontal line indicating the median. Whiskers extend to the most extreme data points within 1.5 × IQR from the lower and upper quartiles. Mean values are shown as dots.

Supplementary Table S12. Statistical analysis of the data distribution for antibiofilm activity of rosemary essential oil (REO) or thyme essential oil (TEO) against *S. aureus* strains (n=10) cultured on different surfaces: agar (A) or biocellulose (BC), and in different media: tryptic soy broth (TSB) or in in vitro wound milieu (IVWM), assessed with an antibiofilm activity of volatile compounds method. Data was grouped by the following growth conditions: surface and medium, or medium only, or surface only. Normal distribution was considered for values of p>0.05. SD- standard deviation, IQR- interquartile range, N- data points, NaN, NA- not applicable. Statistics omitted as all results have identical values.

| Grouping by | Condition | Mean | Median | SD | Min | Max | IQR | N | Skewness | Kurtosis | Shapiro-Wilk p value |
|---|---|---|---|---|---|---|---|---|---|---|---|
| Medium & Surface | TSB/A (REO) | −72.40 | −63.38 | 64.58 | −220.08 | 37.91 | 105.13 | 55 | −0.47 | −0.61 | 0.05 |
| Medium & Surface | IVWM/A (REO) | 11.79 | 9.27 | 47.72 | −78.40 | 100.00 | 63.80 | 59 | 0.11 | −0.64 | 0.17 |
| Medium & Surface | TSB/BC (REO) | −45.94 | −42.08 | 36.38 | −131.19 | 18.63 | 48.05 | 54 | −0.53 | −0.32 | 0.11 |
| Medium & Surface | IVWM/BC (REO) | 8.78 | 6.22 | 43.02 | −108.61 | 97.07 | 64.29 | 60 | −0.34 | 0.08 | 0.46 |
| Medium & Surface | TSB/A (TEO) | 88.39 | 100.00 | 15.48 | 52.40 | 100.00 | 23.78 | 54 | −0.90 | −0.79 | 0.00 |
| Medium & Surface | IVWM/A (TEO) | 100.00 | 100.00 | 0.00 | 100.00 | 100.00 | 0.00 | 60 | NaN | NaN | NA |
| Medium & Surface | TSB/BC (TEO) | 94.61 | 100.00 | 8.32 | 71.52 | 100.00 | 10.04 | 54 | −1.37 | 0.63 | 0.00 |
| Medium & Surface | IVWM/BC (TEO) | 100.00 | 100.00 | 0.00 | 100.00 | 100.00 | 0.00 | 60 | NaN | NaN | NA |
| Medium | TSB (REO) | −58.37 | −47.35 | 51.54 | −194.37 | 37.91 | 71.09 | 109 | −0.64 | −0.26 | 0.00 |
| Medium | IVWM (REO) | 10.27 | 6.82 | 45.25 | −108.61 | 100.00 | 65.06 | 119 | −0.07 | −0.25 | 0.33 |
| Medium | TSB (TEO) | 92.29 | 100.00 | 11.57 | 61.26 | 100.00 | 12.88 | 106 | −1.30 | 0.32 | 0.00 |
| Medium | IVWM (TEO) | 100.00 | 100.00 | 0.00 | 100.00 | 100.00 | 0.00 | 120 | NaN | NaN | NA |
| Surface | A (REO) | −24.82 | −19.46 | 64.74 | −189.57 | 100.00 | 87.53 | 112 | −0.25 | −0.38 | 0.08 |
| Surface | BC (REO) | −17.14 | −17.82 | 48.38 | −131.19 | 97.07 | 57.88 | 114 | −0.06 | −0.23 | 0.70 |
| Surface | A (TEO) | 100.00 | 100.00 | 0.00 | 100.00 | 100.00 | 0.00 | 88 | NaN | NaN | NA |
| Surface | BC (TEO) | 100.00 | 100.00 | 0.00 | 100.00 | 100.00 | 0.00 | 90 | NaN | NaN | NA |

Supplementary Table S13. Parameters of the tested statistic of the differences for antibiofilm activity of rosemary essential oil against *S. aureus* strains (n=10) cultured on different surfaces: agar (A) or biocellulose (BC), and in different media: tryptic soy broth (TSB) or in in vitro wound milieu (IVWM), assessed with an antibiofilm activity of volatile compounds method. Data was grouped by the following growth conditions: surface and medium, or medium only, or surface only. Kruskal-Wallis test, followed by the Dunn's test, was performed. Adjusted p value includes Bonferroni correction. Only differences classified as statistically significant in ANOVA were included. Values of $p<0.05$ were considered significant, $p \leq 0.0001$ was marked with four asterisks. N- data points, ns- no significant differences assessed in post-hoc tests.

| Groupped by | Growth condition 1 | Growth condition 2 | N1 | N2 | mean ranks difference | mean rank1 | mean rank2 | p value | adj. p value | Significance |
|---|---|---|---|---|---|---|---|---|---|---|
| Medium & Surface | IVWM/A | IVWM/BC | 59 | 60 | 0.72 | 151.86 | 152.58 | $9.53 \times 10^{-01}$ | 1.00 | ns |
| Medium & Surface | IVWM/A | TSB/A | 59 | 55 | -85.41 | 151.86 | 66.45 | $4.91 \times 10^{-12}$ | $2.94 \times 10^{-11}$ | **** |
| Medium & Surface | IVWM/A | TSB/BC | 59 | 54 | -71.57 | 151.86 | 80.30 | $8.35 \times 10^{-09}$ | $5.01 \times 10^{-08}$ | **** |
| Medium & Surface | IVWM/BC | TSB/A | 60 | 55 | -86.13 | 152.58 | 66.45 | $2.66 \times 10^{-12}$ | $1.60 \times 10^{-11}$ | **** |
| Medium & Surface | IVWM/BC | TSB/BC | 60 | 54 | -72.29 | 152.58 | 80.30 | $5.15 \times 10^{-09}$ | $3.09 \times 10^{-08}$ | **** |
| Medium & Surface | TSB/A | TSB/BC | 55 | 54 | 13.84 | 66.45 | 80.30 | $2.73 \times 10^{-01}$ | 1.00 | ns |
| Medium | IVWM | TSB | 119 | 109 | -78.91 | 152.23 | 73.31 | $1.82 \times 10^{-19}$ | $1.82 \times 10^{-19}$ | **** |

Supplementary Table S14. Parameters of the tested statistic of the differences for antibiofilm activity of thyme essential oil against *S. aureus* strains (n=10) cultured on different surfaces: agar (A) or biocellulose (BC), and in different media: tryptic soy broth (TSB) or in in vitro wound milieu (IVWM), assessed with an antibiofilm activity of volatile compounds method. Data was grouped by the following growth conditions: surface and medium, or medium only, or surface only. Kruskal-Wallis test, followed by the Dunn's test, was performed. Adjusted p value includes Bonferroni correction. Only differences classified as statistically significant in ANOVA were included. Values of $p<0.05$ were considered significant, $p \leq 0.0001$ was marked with four asterisks. N- data points, ns- no significant differences assessed in post-hoc tests.

| Groupped by | Growth condition 1 | Growth condition 2 | N1 | N2 | mean ranks difference | mean rank1 | mean rank2 | p value | adj. p value | Significance |
|---|---|---|---|---|---|---|---|---|---|---|
| Medium & Surface | IVWM/A | IVWM/BC | 60 | 60 | 0.00 | 139.50 | 139.50 | 1.00 | 1.00 | ns |
| Medium & Surface | IVWM/A | TSB/A | 60 | 54 | -57.67 | 139.50 | 81.83 | $1.22 \times 10^{-10}$ | $7.29 \times 10^{-10}$ | **** |
| Medium & Surface | IVWM/A | TSB/BC | 60 | 54 | -47.89 | 139.50 | 91.61 | $9.00 \times 10^{-08}$ | $5.40 \times 10^{-07}$ | **** |
| Medium & Surface | IVWM/BC | TSB/A | 60 | 54 | -57.67 | 139.50 | 81.83 | $1.22 \times 10^{-10}$ | $7.29 \times 10^{-10}$ | **** |
| Medium & Surface | IVWM/BC | TSB/BC | 60 | 54 | -47.89 | 139.50 | 91.61 | $9.00 \times 10^{-08}$ | $5.40 \times 10^{-07}$ | **** |
| Medium & Surface | TSB/A | TSB/BC | 54 | 54 | 9.78 | 81.83 | 91.61 | $2.87 \times 10^{-01}$ | 1.00 | ns |
| Medium | IVWM | TSB | 120 | 106 | -51.17 | 137.50 | 86.33 | $2.21 \times 10^{-16}$ | $2.21 \times 10^{-16}$ | **** |

Supplementary Table S15. Parameters of the tested statistic of the differences between antibiofilm activity of rosemary essential oil (REO) and thyme essential oil (TEO) against *S. aureus* strains (n=10) cultured on different surfaces: agar (A) or biocellulose (BC), and in different media: tryptic soy broth (TSB) or in in vitro wound milieu (IVWM), assessed with an antibiofilm activity of volatile compounds method. Data was grouped by the following growth conditions: surface and medium, or medium only, or surface only. Kruskal-Wallis test, followed by the Dunn's test, was performed. Adjusted p value includes Bonferroni correction. Only differences classified as statistically significant in ANOVA were included. Values of $p<0.05$ were considered significant, $p \leq 0.0001$ was marked with four asterisks. N- data points.

| Groupped by | Growth condition | Essential oil1 | Essential oil2 | N1 | N2 | mean ranks difference | mean rank1 | mean rank2 | p value | adj. p value | Significance |
|---|---|---|---|---|---|---|---|---|---|---|---|
| Surface | A | REO | TEO | 112 | 88 | 95.54 | 58.46 | 154.00 | $2.50 \times 10^{-34}$ | $2.50 \times 10^{-34}$ | **** |
| Surface | BC | REO | TEO | 114 | 90 | 102.00 | 57.50 | 159.50 | $1.33 \times 10^{-37}$ | $1.33 \times 10^{-37}$ | **** |

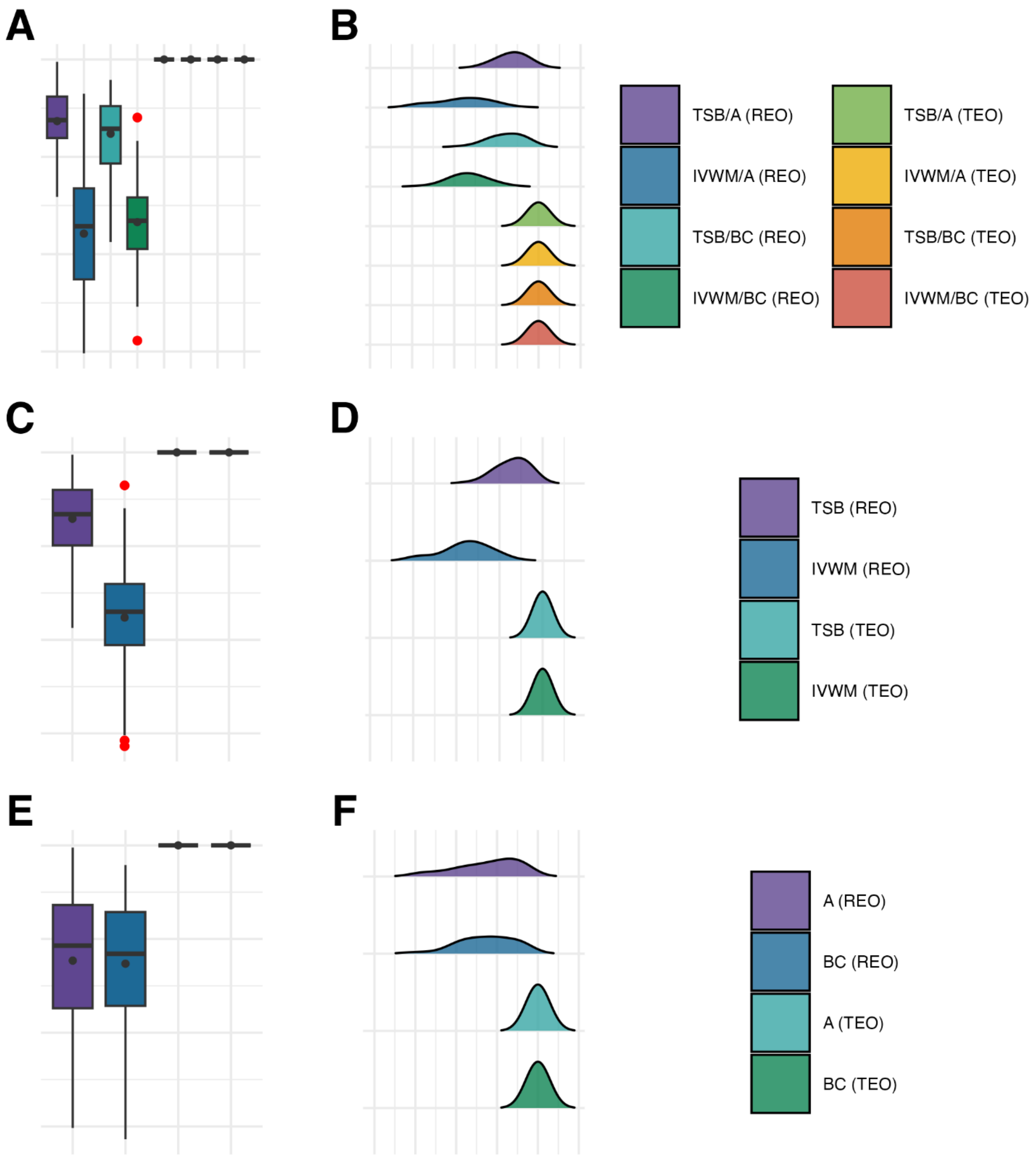


Supplementary Figure S6. Visual representation of the data distribution for antibiofilm activity of rosemary essential oil (REO) or thyme essential oil (TEO) against *S. aureus* strains (n=10) cultured on different surfaces: agar (A) or biocellulose (BC) and in different media: tryptic soy broth (TSB) or in in vitro wound milieu (IVWM), assessed with an antibiofilm dressing's activity measurement method. **(A, B)** Dividing condition: medium and surface. **(C, D)** Dividing condition: medium. **(E, F)** Dividing condition: surface. Each box displays the interquartile range (IQR; 25th to 75th percentiles), with the bold horizontal line indicating the median. Whiskers extend to the most extreme data points within 1.5 × IQR from the lower and upper quartiles. Mean values are shown as dots.

Supplementary Table S16. Statistical analysis of the data distribution for antibiofilm activity of rosemary essential oil (REO) or thyme essential oil (TEO) against *S. aureus* strains (n=10) cultured on different surfaces: agar (A) or biocellulose (BC), and in different media: tryptic soy broth (TSB) or in in vitro wound milieu (IVWM), assessed with an antibiofilm dressing's activity measurement method. Data was grouped by the following growth conditions: surface and medium, or medium only, or surface only. Normal distribution was considered for values of p>0.05. SD- standard deviation, IQR- interquartile range, N- data points, NaN, NA- not applicable. Statistics omitted as all results have identical values.

| Grouping by | Condition | Mean | Median | SD | Min | Max | IQR | N | Skewness | Kurtosis | Shapiro-Wilk p value |
|---|---|---|---|---|---|---|---|---|---|---|---|
| Medium & Surface | TSB/A (REO) | 36.72 | 37.72 | 31.46 | −41.14 | 97.53 | 42.47 | 54 | −0.40 | −0.61 | 0.25 |
| Medium & Surface | IVWM/A (REO) | −78.86 | −71.34 | 64.38 | −201.73 | 64.79 | 93.55 | 57 | −0.18 | −0.73 | 0.31 |
| Medium & Surface | TSB/BC (REO) | 23.75 | 28.85 | 39.21 | −87.38 | 79.01 | 59.04 | 57 | −0.56 | −0.26 | 0.02 |
| Medium & Surface | IVWM/BC (REO) | −67.02 | −65.64 | 43.70 | −188.77 | 40.36 | 52.98 | 57 | −0.06 | 0.12 | 0.98 |
| Medium & Surface | TSB/A (TEO) | 100.00 | 100.00 | 0.00 | 100.00 | 100.00 | 0.00 | 60 | NaN | NaN | NA |
| Medium & Surface | IVWM/A (TEO) | 100.00 | 100.00 | 0.00 | 100.00 | 100.00 | 0.00 | 60 | NaN | NaN | NA |
| Medium & Surface | TSB/BC (TEO) | 100.00 | 100.00 | 0.00 | 100.00 | 100.00 | 0.00 | 60 | NaN | NaN | NA |
| Medium & Surface | IVWM/BC (TEO) | 100.00 | 100.00 | 0.00 | 100.00 | 100.00 | 0.00 | 60 | NaN | NaN | NA |
| Medium | TSB (REO) | 29.18 | 34.03 | 37.09 | −87.38 | 97.53 | 59.35 | 112 | −0.65 | 0.02 | 0.00 |
| Medium | IVWM (REO) | −76.29 | −70.09 | 58.23 | −213.61 | 64.79 | 65.27 | 117 | −0.35 | −0.20 | 0.08 |
| Medium | TSB (TEO) | 100.00 | 100.00 | 0.00 | 100.00 | 100.00 | 0.00 | 120 | NaN | NaN | NA |
| Medium | IVWM (TEO) | 100.00 | 100.00 | 0.00 | 100.00 | 100.00 | 0.00 | 120 | NaN | NaN | NA |
| Surface | A (REO) | −23.03 | −7.06 | 76.93 | −201.73 | 97.53 | 110.15 | 112 | −0.62 | −0.57 | 0.00 |
| Surface | BC (REO) | −26.31 | −15.71 | 67.28 | −213.61 | 79.01 | 100.21 | 117 | −0.45 | −0.26 | 0.00 |
| Surface | A (TEO) | 100.00 | 100.00 | 0.00 | 100.00 | 100.00 | 0.00 | 120 | NaN | NaN | NA |
| Surface | BC (TEO) | 100.00 | 100.00 | 0.00 | 100.00 | 100.00 | 0.00 | 120 | NaN | NaN | NA |

Supplementary Table S17. Parameters of the tested statistic of the differences for antibiofilm activity of rosemary essential oil against *S. aureus* strains (n=10) cultured on different surfaces: agar (A) or biocellulose (BC), and in different media: tryptic soy broth (TSB) or in in vitro wound milieu (IVWM), assessed with an antibiofilm dressing's activity measurement method. Data was grouped by the following growth conditions: surface and medium, or medium only, or surface only. Kruskal-Wallis test, followed by the Dunn’s test, was performed. Adjusted p value includes Bonferroni correction. Only differences classified as statistically significant in ANOVA were included. Values of $p<0.05$ were considered significant, $p \leq 0.0001$ was marked with four asterisks. N- data points, ns- no significant differences assessed in post-hoc tests.

| Groupped by | Growth condition 1 | Growth condition 2 | N1 | N2 | mean ranks difference | mean rank1 | mean rank2 | p value | adj. p value | Significance |
|---|---|---|---|---|---|---|---|---|---|---|
| Medium & Surface | IVWM/A | IVWM/BC | 57 | 57 | 2.98 | 62.28 | 65.26 | $8.07 \times 10^{-01}$ | 1.00 | ns |
| Medium & Surface | IVWM/A | TSB/A | 57 | 54 | 108.52 | 62.28 | 170.80 | $1.66 \times 10^{-18}$ | $9.96 \times 10^{-18}$ | **** |
| Medium & Surface | IVWM/A | TSB/BC | 57 | 57 | 94.42 | 62.28 | 156.70 | $9.67 \times 10^{-15}$ | $5.80 \times 10^{-14}$ | **** |
| Medium & Surface | IVWM/BC | TSB/A | 57 | 54 | 105.53 | 65.26 | 170.80 | $1.38 \times 10^{-17}$ | $8.26 \times 10^{-17}$ | **** |
| Medium & Surface | IVWM/BC | TSB/BC | 57 | 57 | 91.44 | 65.26 | 156.70 | $6.44 \times 10^{-14}$ | $3.86 \times 10^{-13}$ | **** |
| Medium & Surface | TSB/A | TSB/BC | 54 | 57 | -14.09 | 170.80 | 156.70 | $2.54 \times 10^{-01}$ | 1.00 | ns |
| Medium | IVWM | TSB | 117 | 112 | 101.04 | 65.58 | 166.62 | $8.56 \times 10^{-31}$ | $8.56 \times 10^{-31}$ | **** |

Supplementary Table S18. Parameters of the tested statistic of the differences between antibiofilm activity of rosemary essential oil (REO) and thyme essential oil (TEO) against *S. aureus* strains (n=10) cultured on different surfaces: agar (A) or biocellulose (BC), and in different media: tryptic soy broth (TSB) or in in vitro wound milieu (IVWM), assessed with an antibiofilm dressing's activity measurement method. Data was grouped by the following growth conditions: surface and medium, or medium only, or surface only. Kruskal-Wallis test, followed by the Dunn’s test, was performed. Adjusted p value includes Bonferroni correction. Only differences classified as statistically significant in ANOVA were included. Values of $p<0.05$ were considered significant, $p \leq 0.0001$ was marked with four asterisks. N- data points.

| Groupped by | Growth condition | Essential oil1 | Essential oil2 | N1 | N2 | mean ranks difference | mean rank1 | mean rank2 | p value | adj. p value | Significance |
|---|---|---|---|---|---|---|---|---|---|---|---|
| Medium & Surface | TSB/A | REO | TEO | 54 | 60 | 57.00 | 27.50 | 84.50 | $2.60 \times 10^{-23}$ | $2.60 \times 10^{-23}$ | **** |
| Medium & Surface | IVWM/A | REO | TEO | 57 | 60 | 58.50 | 29.00 | 87.50 | $1.18 \times 10^{-23}$ | $1.18 \times 10^{-23}$ | **** |
| Medium & Surface | TSB/BC | REO | TEO | 57 | 60 | 58.50 | 29.00 | 87.50 | $1.18 \times 10^{-23}$ | $1.18 \times 10^{-23}$ | **** |
| Medium & Surface | IVWM/BC | REO | TEO | 57 | 60 | 58.50 | 29.00 | 87.50 | $1.18 \times 10^{-23}$ | $1.18 \times 10^{-23}$ | **** |

Supplementary Table S19. Parameters of the tested statistic of the differences between the results of two methods (an antibiofilm activity of volatile compounds vs an antibiofilm dressing's activity measurement) assessing the antibiofilm activity of rosemary essential oil (REO) or thyme essential oil (TEO) against *S. aureus* strains (n=10) cultured on different surfaces: agar (A) or biocellulose (BC), and in different media: tryptic soy broth (TSB) or in in vitro wound milieu (IVWM). Data was grouped by the following growth conditions: surface and medium, or medium only, or surface only. Kruskal-Wallis test, followed by the Dunn’s test, was performed. Adjusted p value includes Bonferroni correction. Only differences classified as statistically significant in ANOVA were included. Values of $p<0.05$ were considered significant, $p \leq 0.0001$ was marked with four asterisks. N- data points.

| Groupped by | Essential oil | Growth condition | Method1 | Method2 | N1 | N2 | mean ranks difference | mean rank1 | mean rank2 | p value | adj. p value | Significance |
|---|---|---|---|---|---|---|---|---|---|---|---|---|
| Medium & Surface | REO | TSB/A | ABV | ADAM | 55 | 55 | 48.64 | 31.18 | 79.82 | $1.29 \times 10^{-15}$ | $1.29 \times 10^{-15}$ | **** |
| Medium & Surface | REO | IVWM/A | ABV | ADAM | 60 | 60 | -44.23 | 82.62 | 38.38 | $3.28 \times 10^{-12}$ | $3.28 \times 10^{-12}$ | **** |
| Medium & Surface | REO | TSB/BC | ABV | ADAM | 57 | 57 | 46.58 | 34.21 | 80.79 | $5.35 \times 10^{-14}$ | $5.35 \times 10^{-14}$ | **** |
| Medium & Surface | REO | IVWM/BC | ABV | ADAM | 60 | 60 | -47.93 | 84.47 | 36.53 | $4.44 \times 10^{-14}$ | $4.44 \times 10^{-14}$ | **** |
| Medium & Surface | TEO | TSB/A | ABV | ADAM | 57 | 60 | 29.76 | 43.74 | 73.50 | $3.88 \times 10^{-10}$ | $3.88 \times 10^{-10}$ | **** |
| Medium & Surface | TEO | TSB/BC | ABV | ADAM | 57 | 60 | 27.71 | 44.79 | 72.50 | $2.18 \times 10^{-09}$ | $2.18 \times 10^{-09}$ | **** |
